# Integration of hunger and hormonal state gates infant-directed aggression

**DOI:** 10.1101/2024.11.25.625278

**Authors:** Mingran Cao, Rachida Ammari, Maxwell X. Chen, Patty Wai, Aashna Sahni, Swang Liang, Nathalie Legrave, James Macrae, Molly Strom, Johannes Kohl

## Abstract

Social behaviour is profoundly shaped by internal physiological states. While significant progress has been made in understanding how *individual* states such as hunger, stress, or arousal modulate behaviour, animals experience *multiple* states at any given time. The neural mechanisms that integrate such orthogonal states—and how this integration affects behaviour—remain poorly understood. Here we report how hunger and estrous state converge on neurons in the medial preoptic area (MPOA) to shape infant-directed behaviour. We find that hunger promotes pup-directed aggression in normally non-aggressive virgin female mice. This behavioural switch occurs through inhibition of MPOA neurons, driven by the release of neuropeptide Y (NPY) from Agouti-related peptide-expressing neurons in the arcuate nucleus (Arc^AgRP^ neurons). The propensity for hunger-induced aggression is set by reproductive state, with MPOA neurons detecting changes in progesterone (P4) to estradiol (E2) ratio across the estrous cycle. Hunger and estrous state converge on HCN (hyperpolarization-activated cyclic nucleotide-gated) channels, which sets the baseline activity and excitability of MPOA neurons. Using micro-endoscopic imaging, we confirm these findings *in vivo,* revealing that MPOA neurons encode a state for pup-directed aggression. This work thus provides a mechanistic understanding of how multiple physiological states are integrated to flexibly control social behaviour.

## Main

When encountering conspecifics, animals must decide on how to behave. Such social decisions are typically seen as the result of accumulating external, sensory information about the target (e.g., sex, age, status). However, internal states—such as hunger, stress or arousal—profoundly affect social behaviour. While the effects of individual physiological states on behaviour are increasingly well understood^4–15^, organisms experience multiple states at any given time^3^. How such states are integrated in the brain to shape social behaviour remains largely unknown. We have addressed this question using a simple paradigm in which female mice are faced with pups and exhibit either pup-directed care or aggression. We first establish how two state variables, hunger and estrous state, affect pup interactions, before uncovering the cellular and neural mechanisms by which these orthogonal states are integrated to shape social behaviour.

### Arc^AgRP^→MPOA projections mediate hunger-induced, pup-directed aggression

Virgin female laboratory mice typically either ignore pups or exhibit spontaneous parental behaviour^16^. We found that food deprivation induces a shift toward pup-directed aggression in these animals (Fig. 1a). The percentage of aggressive mice (Agg^+^) increased, and attack latency decreased, with food deprivation duration, plateauing after 3 h (Fig. 1b and Extended Data Fig. 1a). Restoring food access increased feeding and diminished pup-directed aggression (Fig. 1b, c). Importantly, food deprivation triggered aggression, regardless of whether mice had previously shown parental behaviour or ignored pups, with similar attack latencies in both groups (Extended Data Fig. 1b, c). This aggression was specifically directed at pups, as interactions with prey or adult intruders of either sex were unaffected by food deprivation (Extended Data Fig. 1e). Moreover, this behavioural shift was not stress-related; food-deprived mice showed no changes in performance on elevated plus maze and open field tests (Extended Data Fig. 1f, g) and were unaffected by other stressors (Extended Data Fig. 1h). Hunger therefore triggers a switch to infant-directed aggression in virgin female mice.

**Fig. 1:**
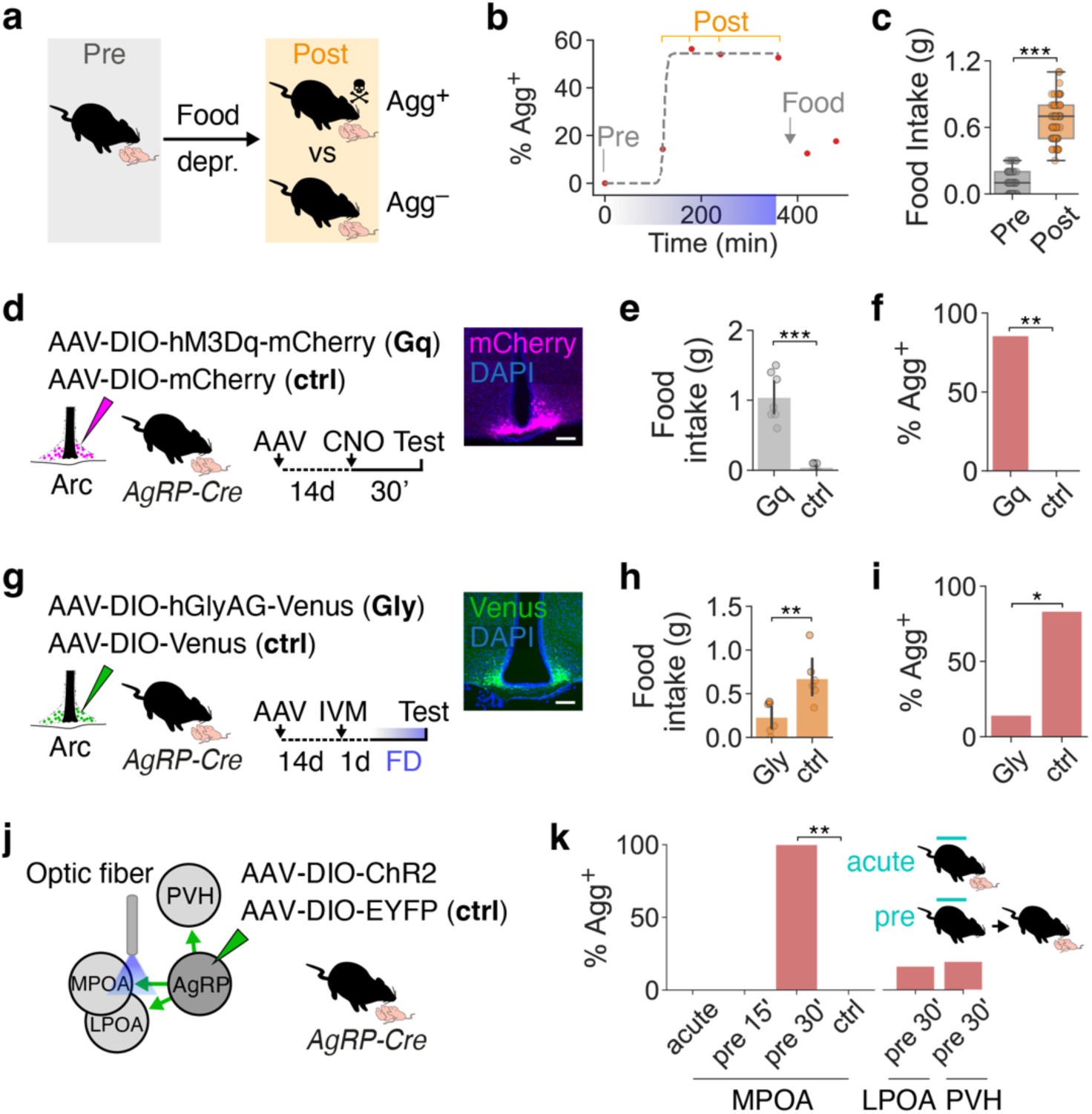
Arc^AgRP^→MPOA projections mediate hunger-induced, pup-directed aggression. **a,** Food deprivation-induced switch to pup-directed aggression in virgin female mice (Pre, before, Post, after food deprivation; Agg^+^, aggressive; Agg^−^, non-aggressive). **b,** Percentage of Agg^+^ mice as a function of food deprivation duration (Re-fed for 1 or 2 h, each point represents a cohort of n = 9–10 mice; blue bar, deprivation period; logistic regression, *r*^2^ = 0.99) **c,** 1-h food intake before and after 6-h food deprivation (n = 40). **d,** Chemogenetic activation of Arc^AgRP^ neurons (Gq) compared to control. Example brain section with mCherry in Arc^AgRP^ neurons. Scale bar, 100 µm. **e,** Effect of chemogenetic Arc^AgRP^ neuron activation on 1-h food consumption in sated mice, and control (n = 7, 7). **f,** Percentage of sated mice injected with CNO showing aggression, and control (n = 7, 7). **g,** Chemogenetic inhibition of Arc^AgRP^ neurons with ivermectin (IVM)-sensitive Gly receptor vs control after 6-h food deprivation. **h,** Effect of chemogenetic Arc^AgRP^ neuron inhibition on 1-h food consumption in food-deprived mice, and control (n = 6, 6). **i,** Percentage of food-deprived mice injected with IVM showing aggression, and controls (n = 7, 6). **j,** Optogenetic activation of Arc^AgRP^ candidate projections. **k**, Percentage of mice exhibiting pup-directed aggression after optogenetic stimulation of Arc^AgRP^ projections. Stimulation during (acute), or for 15 or 30 min before (pre) pup interactions (n = 5, 6, 5, 6 for MPOA, LPOA, PVH, ctrl). Control mice received 30-min pre-stimulation. Paired *t* test in **c**. *U* test in **e** and **h**. Fisher’s exact test in **f, i** and **k** (Benjamini-Hochberg adjustment in **k**). Data are mean ± s.e.m. Box plots show median, 25^th^, 75^th^ percentiles, and whiskers show range. \*\*\**P* < 0.001; \*\**P* < 0.01; \**P* < 0.05.

We next investigated the neural mechanisms underlying this switch. Arc^AgRP^ neurons play a central role in regulating hunger-driven behaviours^17,18^. We therefore tested whether they mediate the effects of food deprivation on pup-directed behaviour. Chemogenetic activation of Arc^AgRP^ neurons increased food consumption, as previously reported (Fig. 1d, e)^18,19^. Strikingly, this manipulation also elicited pup-directed aggression in sated mice, whereas no effects were observed in animals injected with a control virus (Fig. 1d–f). Activation of Arc^AgRP^ neurons is therefore sufficient to evoke pup-directed aggression in females. Conversely, when Arc^AgRP^ neurons were chemogenetically inhibited via an ivermectin-responsive human glycine receptor (hGlyR)^20,21^, food deprivation-evoked pup aggression was strongly diminished (Fig. 1g–i).

To address whether hungry mice attack pups because they perceive them as food, we recorded bulk Arc^AgRP^ activity using fiber photometry (Extended Data Fig. 1i)^22^. Arc^AgRP^ activity is known to increase during food deprivation and to quickly decrease in response to food-related cues^23,24^. In contrast, we observed that it increased during pup investigation (Extended Data Fig. 1j, k), similar to previously reported responses to adult conspecifics^25^. This suggests that aggressive females do not perceive pups as food.

Arc^AgRP^ neurons might exert these effects by directly targeting circuits that mediate pup-directed behaviour^26,27^. We used the immediate early gene *c-fos* as an indirect readout of neuronal recruitment between aggressive (Agg^+^) and non-aggressive (Agg^−^) mice (Extended Data Fig. 2a, b). We focused our analysis on the hypothalamus, which contains neurons critical for pup-directed behaviours^26,28,29^. c-Fos^+^ cell densities significantly differed between Agg^+^ and Agg^−^ mice in 5 out of the 53 assessed brain areas (Extended Data Fig. 2c). All 5 areas exhibited lower c-Fos^+^ cell densities in Agg^+^ mice, suggesting that Arc^AgRP^ neurons might drive pup-directed aggression by inhibiting parenting circuits. Arc^AgRP^ neurons send largely non-collateralised projections to more than a dozen downstream areas^13,30–34^, including to 2 of these 5 candidate areas: the MPOA and the lateral preoptic area (LPOA) (Extended Data Fig. 2d)^35–37^. To address whether these candidate projections mediate pup-directed aggression, we virally expressed channelrhodopsin-2 (ChR2) in Arc^AgRP^ neurons and implanted optical fibers above their projection targets (Fig. 1j and Extended Data Fig. 3a). Acute optogenetic stimulation of Arc^AgRP^→MPOA projections during pup interactions, or 15 min of stimulation prior to behavioural testing, did not affect pup-directed behaviour (Fig. 1k). Strikingly, however, stimulating MPOA projections for 30 min before pup interactions switched all sated mice to pup-directed aggression (Fig. 1k). Prolonged stimulation of MPOA projections for 1 h (see ^38^) also increased food intake; this increase, however, was correlated with longer attack latencies (Extended Data Fig. 3b–e), indicating that Arc^AgRP^→MPOA projections influence feeding and pup-directed aggression via dissociable mechanisms. In contrast, optogenetic stimulation of nearby LPOA projections did not affect pup-directed behaviour or food intake (Fig. 1k and Extended Data Fig. 3g). We also confirmed that activation of projections to the paraventricular nucleus of the hypothalamus (PVH) increased food intake, as previously shown^39^, without affecting social behaviour (Fig. 1k and Extended Data Fig. 3h). Arc^AgRP^→MPOA projections thus mediate hunger-induced pup aggression.

### Estrous state sets switching probability in the MPOA

We next asked why hunger elicits pup-directed aggression in only ∼50% of females (Fig. 1b). Agg^+^ mice were not hungrier than Agg^−^ mice because food consumption and plasma levels of the hunger hormone ghrelin did not significantly differ between the two groups (Fig. 2a–c). We therefore hypothesised that Agg^+^ females are in a reproductive state permissive to aggression. In female rodents, the estrous cycle lasts 4–5 days and is linked with pronounced behavioural and neurophysiological changes (Fig. 2d)^40–44^. The percentage of mice switching to pup-directed aggression (switching rate) fluctuated across estrous cycle stages, being highest in metestrus (70%) and lowest in estrus (32%, Fig. 2e). Estradiol (E2) and progesterone (P4) are main effectors of the estrous cycle (Fig. 2d)^45^, but switching rate was not correlated with individual levels of E2 or P4 (Extended Figure 4a, b). Instead, it tracked 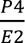 ratio, suggesting that relative levels of both hormones are integrated in feeding and/or parenting circuits (Fig. 2f).

**Fig. 2:**
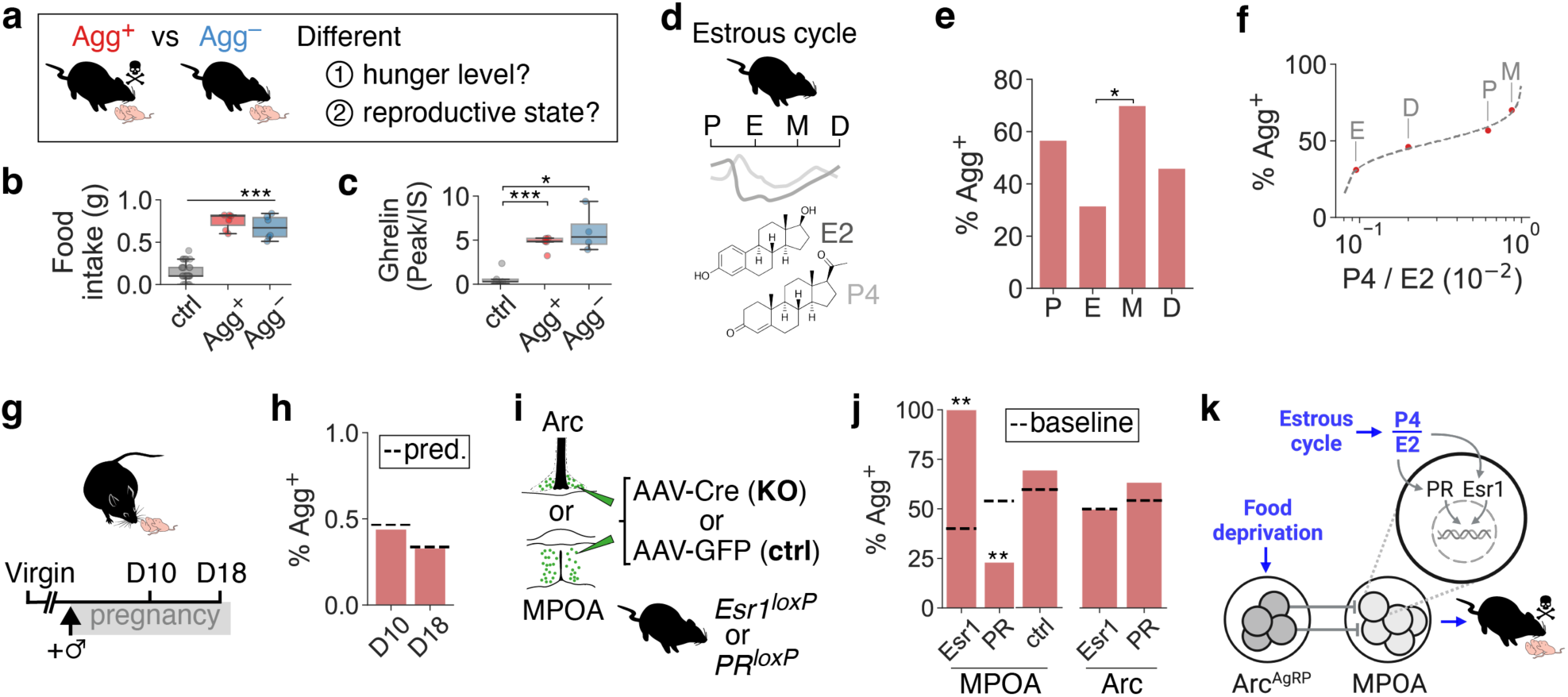
Estrous state sets behavioural switching probability in the MPOA. **a,** Potential factors contributing to different pup-directed behaviours after food deprivation. **b,** 1-h food intake at baseline (ctrl) and after 6-h food deprivation. **c,** Relative plasma ghrelin levels (ratio of peak area to internal standard) at baseline (ctrl) and after 6-h food deprivation. **d,** Plasma E2 and P4 levels across estrous cycle (P, proestrus, E, estrus, M, metestrus, D, diestrus). **e,** Percentage of Agg^+^ mice across estrous stages (n = 30, 19, 40, 37 mice). **f**, Switching rate as a function of 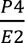 ratio (logit, *r*^2^ = 0.99, n = 30, 19, 40, 37 mice; Extended Data Table 2). **g**, Testing at mid-(day 10, D10) and late (D18) pregnancy. **h,** Switching rates in pregnant females. Dashed lines indicate predicted values (see Methods). **i,** Targeted Esr1 or PR ablation in MPOA or Arc, and control. **j**, Percentage of Agg^+^ mice after Esr1 or PR KO (n = 10, 13, 8, 10, 11 mice). Dashed lines indicate predicted baseline switching rates of each cohort with intact receptors (see Methods). **k**, Integration of estrous and hunger state in the MPOA sets switching rate. One-way ANOVA with Tukey *post hoc* test in **b, c**. Fisher’s exact test with Benjamini-Hochberg adjustment in **e.** Poisson binomial distribution for expected aggression rates based on estrous state in **h, j**; observed rates outside the 99% confidence interval are indicated.

**Fig. 3:**
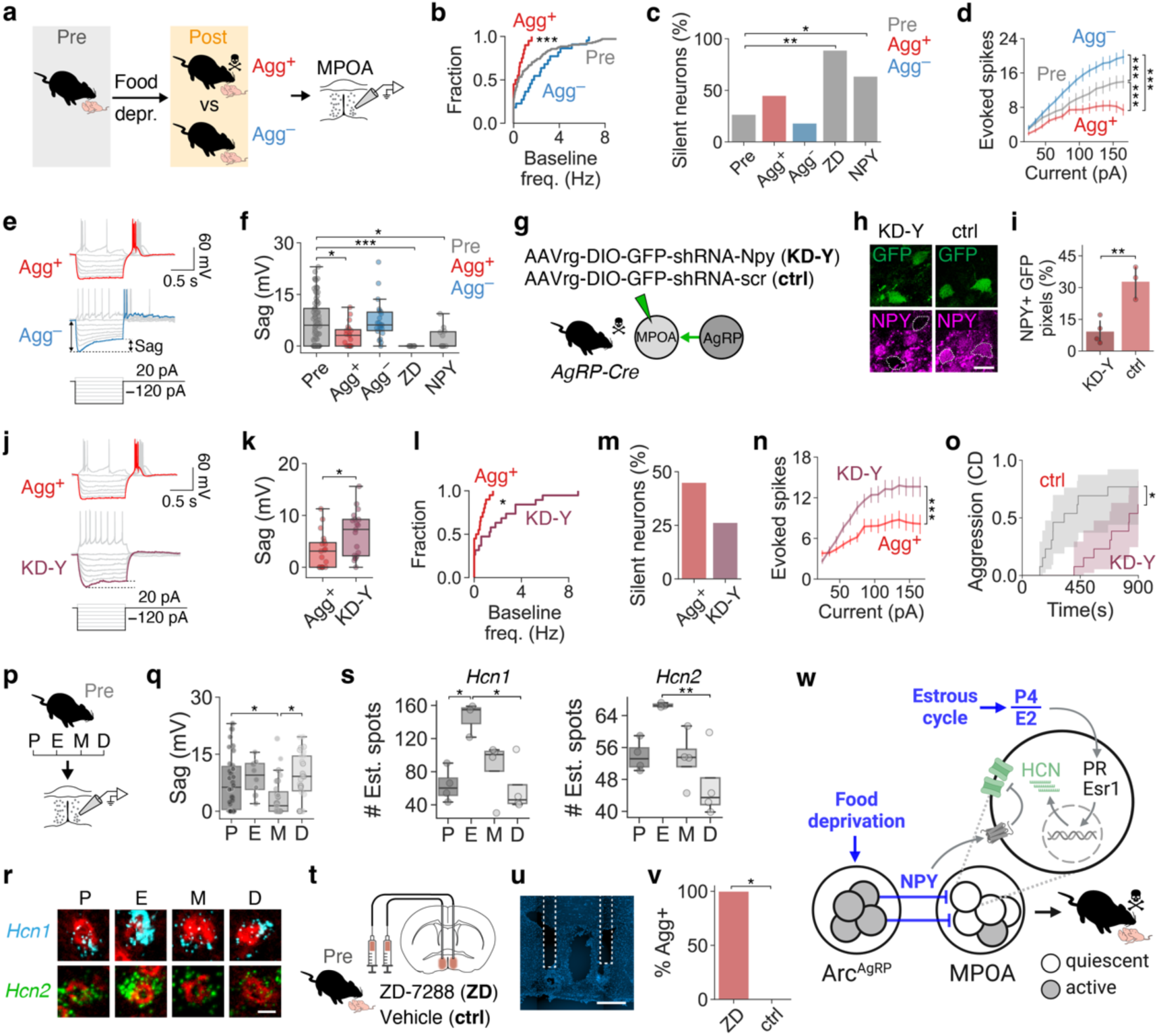
Integration of hunger and estrous state in MPOA neurons. **a,** Whole-cell recordings from MPOA neurons. **b,** Baseline firing (Pre, Agg^+^, Agg^−^: 105, 20, 22 cells from n = 17, 6, 3 mice). **c,** Percentage of silent neurons at resting membrane potential (105, 20, 22, 9, 11 cells from n = 17, 6, 3, 1, 4 mice). **d,** Action potentials per injected current (Pre, Agg^+^, Agg^−^:23, 22, 21 cells from n = 7, 5, 3 mice). **e**, Example current clamp recordings with voltage sag highlighted. **f,** Voltage sag amplitude (82, 19, 22, 9, 11 cells from n = 15, 6, 3, 1, 4 mice). **g,** NPY knockdown (KD-Y) in Arc^AgRP^→MPOA projections. **h,** Example images of NPY knockdown and control. Scale bar, 20 µm. **i**, NPY knockdown efficiency (n = 4, 3 mice). **j,** Current clamp recordings from MPOA neurons in Agg^+^ and Agg^+^ KD-Y mice. **k,** Voltage sag amplitude (19, 19 cells from n = 6, 3 mice). **l,** Baseline firing (20, 19 cells from 6, 3 mice). **m,** Percentage of silent neurons at resting membrane potential (20, 19 cells from n = 6, 3 mice). **n,** Action potentials per injected current (Agg^+^, KD-Y: 22, 17 cells from n = 5, 3 mice). **o**, Cumulative incidence of aggression (n = 8, 13 mice). Shaded areas are confidence intervals. **p,** MPOA recordings across estrous cycle. **q,** Voltage sag amplitude across estrous cycle (23, 7, 22, 27 neurons from n = 7, 2, 4, 2 mice). **r, s,** *Hcn1* and *Hcn2* mRNA expression across estrous cycle (**r,** red, NeuN counterstain; scale bars, 10 µm) and quantification (**s,** estimated number of spots, see Methods; n = 4, 3, 4, 4 mice). **t,** Bilateral infusion of HCN blocker (ZD) or vehicle (ctrl) into the MPOA. **u,** Cannula placement in MPOA. Scale bar, 1 mm. **v,** Percentage of Agg^+^ mice after ZD application, and control (n = 3, 5). **w,** Integration of hunger and estrous state in MPOA neurons. *U* test in **b, i, k, l** (Pre vs Agg^+^ in **b**). Fisher’s exact test in **c, m, v** (Benjamini-Hochberg adjustment in **c**). Two-way ANOVA with Tukey *post hoc* test in **d, n**. *U* test with Benjamini-Hochberg adjustment (Pre, Agg^+^, Agg^−^; Pre, ZD, NPY) in **f**. Log-rank test in **o**. One-way ANOVA with Tukey *post hoc* test in **q, s**.

**Fig. 4:**
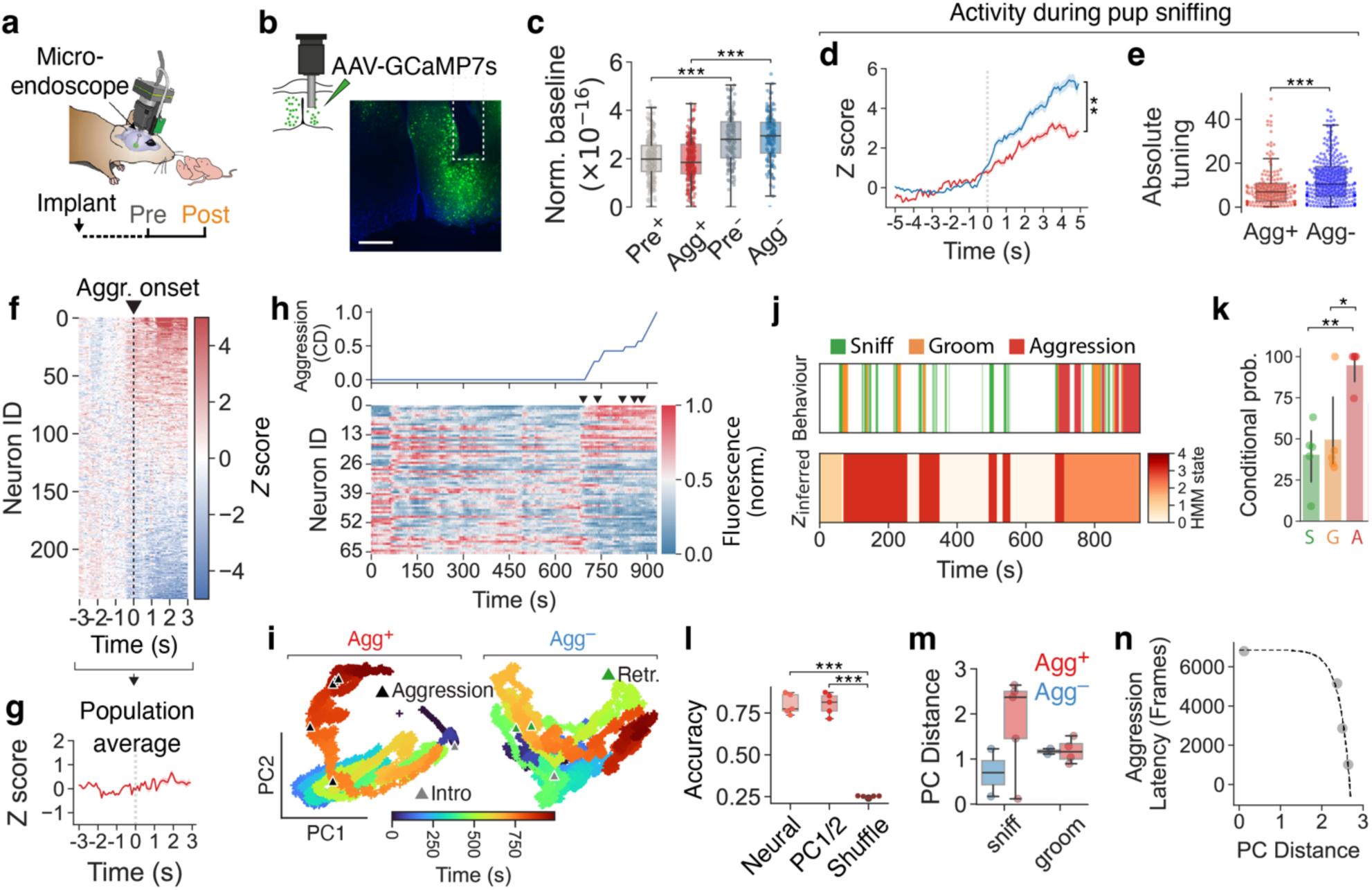
MPOA neurons encode an aggressive state. **a,** Miniature microscope recordings during pup interactions. **b,** GCaMP expression and GRIN lens placement in MPOA. Scale bar, 500 µm. **c,** Normalised neuronal baseline activity (Pre^+^ and Pre^−^ refer to mice classified as Agg^+^ or Agg^−^ after subsequent food deprivation). Raw fluorescence values shown. **d, e,** Averaged, *Z*-scored responses (**d**) and absolute tuning index (**e**) of MPOA neurons during pup chemoinvestigation (114, 106 neurons from n = 4, 2 mice). **f,** Temporal profile of neuronal responses during pup-directed aggression with hierarchical clustering based on mean response onset (243 neurons from n = 5 mice). **g,** Averaged, *Z*-scored responses during aggression onset. **h,** Example heatmap of neuronal responses from single Agg^+^ animal, sorted based on correlation of activity with cumulative distribution of aggression (n = 67 neurons). Arrows indicate aggression episodes. **i,** Population activity traces projected onto first two PCs. Note the Agg^+^-specific state along PC2. **j,** Example HMM state segmentation using Agg^+^ neural data *(bottom)*, and ethogram *(top)*. **k,** Conditional probability of observing the indicated behaviours when the system is in the Hidden Markov Model (HMM) state that most frequently aligns with those behaviours (S, sniff, G, groom, A, aggression; n = 5, 5, 5 mice). **l,** Behavioural prediction accuracy from multiclass SVM classifier trained on neural data, PC1 and PC2, or shuffled data (n = 5 mice). **m,** Pre vs Post PC distances during pup chemoinvestigation and grooming (see Methods, n = 5, 2 mice). **n,** Exponential fit of PC distance vs aggression latency in Agg^+^ mice (n = 4). Mixed linear model with mouse ID as group in **c**. *U* test in **d, e**. One-way ANOVA with Tukey *post hoc* test in **k**, **l, m**.

Supporting this hypothesis, the switching rates of females in mid- or late pregnancy—which have higher P4 and E2 levels than virgins, but comparable 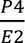 ratios (Fig. 2g and Extended Data Table 2)^46,47^—closely matched our predictions (Fig. 2h). In contrast, estrous state did not affect baseline pup-directed behaviour or attack latency (Extended Data Fig. 4c, d).

We next tested this model and assessed where hormonal state is sensed in this context. E2 and P4 can influence neuronal function both through membrane-bound receptors and through their intracellular receptors, Esr1 and PR, which act as transcription factors^48–51^ and are highly enriched in the MPOA^52^. Mice carrying floxed Esr1 or PR alleles were injected into the Arc or MPOA with an adeno-associated virus (AAV) expressing Cre recombinase (Fig. 2i)^7,53^. This resulted in local receptor knockout (KO), whereas injection of a control AAV did not affect receptor expression^7^. KO of Esr1 or PR in the Arc did not alter pup-directed behaviour, but receptor ablation in the MPOA significantly affected switching rate: 100% of Esr1-ablated mice became aggressive after food deprivation, compared to a predicted 40% baseline rate for mice with intact receptors (Fig. 2j, see Methods). This likely occurs because E2-insensitivity increases the relative 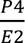 ratio sensed by MPOA neurons. In contrast, only 23% of PR-ablated mice became aggressive (53% predicted), in accordance with a low 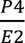 ratio acting on MPOA neurons (Fig. 2j). Levels of parental behaviour were positively correlated with PR ablation efficiency (Extended Data Fig. 4g), and injection of a control AAV did not affect switching rate (Fig. 2j). Importantly, food intake was not affected by estrous state, supporting the conclusion that ovarian hormones modulate parenting rather than feeding centres in this context (Extended Data Fig. 4e, f). Integration of hunger and estrous state in the MPOA thus controls a switch in pup-directed behaviour (Fig. 2k).

### Integration of hunger and estrous state in MPOA neurons

To address how MPOA neurons perform this integration, we performed patch clamp recordings in brain slices from female mice before (Pre) and after food deprivation (Fig. 3a). MPOA neurons from Agg^+^ mice exhibited reduced spontaneous firing, a twofold increase in the proportion of silent neurons, and a strong reduction in intrinsic excitability (Fig. 3b–d and Extended Data Fig. 5a–c). Other biophysical parameters were unchanged (Extended Data Fig. 5f–r). These changes also occurred in Galanin (Gal)-expressing MPOA neurons, which have a well-defined role in parental behaviour (Extended Data Fig. 6)^54,55^. The reduced spontaneous activity and excitability of MPOA neurons in Agg^+^ mice were not due to changes in resting membrane potential or synaptic inputs (Extended Data Fig. 5f, m–p). However, membrane input resistance was increased in Agg^+^ mice, hinting at closure or downregulation of ion channels (Extended Data Fig. 5d). Indeed, negative current injection revealed a depolarising voltage sag in Pre and Agg^−^ mice which was strongly reduced in Agg^+^ mice (Fig. 3e, f) and inversely correlated with input resistance (Extended Data Fig. 5e). This sag is mediated by hyperpolarization-activated cyclic nucleotide–gated (HCN) channels and was abolished by the HCN blocker ZD-7288 (ZD) (Fig. 3f)^56,57^. HCN blockade in brain slices from sated mice also silenced MPOA neurons (Fig. 3c and Extended Data Fig. 8a) and decreased their excitability (Extended Data Fig. 8d). Interfering with HCN channel function therefore reproduces the Agg^+^ neuronal phenotype in the MPOA.

In addition to the neuropeptide AgRP itself, Arc^AgRP^ neurons release γ-aminobutyric acid (GABA) and Neuropeptide Y (NPY), which mediate feeding in a partially redundant manner^19,58–60^. We thus asked which of these neuromediators control the effect of food deprivation on pup-directed behaviour. Food deprivation affects pup interactions within ∼2 h (Fig. 1b) and optogenetic activation of Arc^AgRP^→MPOA projections for 30 min results in pup-directed aggression (Fig. 1k). In contrast, AgRP mediates a delayed, chronic feeding response^19,61^ and its application to brain slices from sated mice did not reproduce the MPOA neuronal silencing observed in Agg^+^ animals (Extended Data Fig. 7a–f). GABA and NPY modulate feeding more rapidly^19,61^, but we found no evidence for direct GABAergic Arc^AgRP^→MPOA connectivity (Extended Data Fig. 7m–p and Supplementary Note). In addition, food deprivation did not hyperpolarise MPOA neurons, as expected from increased GABAergic transmission (Extended Data Fig. 5f). We therefore reasoned that NPY release from Arc^AgRP^→MPOA projections during food deprivation mediates hunger-evoked aggression. Indeed, bath application of NPY to brain slices from sated animals partially reproduced the neural phenotype of Agg^+^ mice, reducing MPOA neuronal activity and the HCN-mediated voltage sag (Fig. 3c, f and Extended Data Fig. 7g–l). To directly test the role of NPY release from Arc^AgRP^→MPOA projections, we injected a retrograde, Cre-dependent AAV expressing a short hairpin RNA (shRNA) against NPY into the MPOA of *AgRP-Cre* mice (Fig. 3g). This led to NPY knockdown (KD-Y) in Arc^AgRP^→MPOA projections, whereas a control virus did not affect NPY expression (Fig. 3g–i). Projection-specific NPY knockdown in Agg^+^ mice increased sag amplitude (i.e., HCN function), reduced neuronal silencing, and increased excitability (Fig. 3j–n). Strikingly, this manipulation delayed the onset of pup-directed aggression (Fig. 3o), with attack latency showing a correlation to the number of transduced neurons (Extended Data Fig. 9a). In contrast, NPY knockdown did not affect food intake after food deprivation in Agg^+^ and Agg^−^ mice (Extended Data Fig. 9b). NPY release from Arc^AgRP^→MPOA projections therefore promotes the hunger-evoked switch to aggression.

How does reproductive state affect this system to set switching rate? Our receptor KO experiments suggest that estrous state is sensed in the MPOA (Fig. 2i, j). We thus performed whole-cell recordings from MPOA neurons of females across estrous stages (Fig. 3p and Extended Data Fig. 10). Voltage sag amplitude and the proportion of neurons exhibiting voltage sag fluctuated during the estrous cycle (Fig. 3q and Extended Data Fig. 10f), being lowest in metestrus—when switching rate is maximal—and highest in estrus, when switching rate is minimal (Fig. 2f). Switching rate was also inversely correlated with the proportion of sag-exhibiting neurons (Extended Data Fig. 10g). We therefore hypothesised that fluctuating 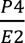 ratio tunes HCN expression in MPOA neurons via the transcription factor receptors PR and Esr1. HCN channels are comprised of 4 subunits (HCN1–4), all of which are expressed in the MPOA (Extended Data Fig. 11a, b)^62–65^. Using single-molecule fluorescence *in situ* hybridization, we found that *Hcn1, Hcn2* and *Hcn4* transcript levels in MPOA neurons fluctuated across the estrous cycle, with *Hcn1* and *Hcn2* showing a pronounced peak in estrus (Fig. 3r, s and Extended Data Fig. 11c, d).

Hunger and estrous state thus converge on HCN channels to regulate MPOA neuronal activity and excitability. While estrous stage sets HCN channel number, food deprivation inhibits available HCN channels via NPY signalling. In estrus, a low 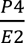 ratio results in a high density of HCN channels, which are only partially inhibited by NPY. As a result, MPOA neurons remain active and excitable even after food deprivation. In contrast, the high 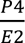 ratio during metestrus reduces HCN channel number, allowing for more effective inhibition by NPY. This leads to quiescent MPOA neurons with low activity and excitability, promoting aggression (Fig. 3w). To test this model, we administered HCN channel blocker into the MPOA of non-food-deprived mice before behavioural testing (Fig. 3t, u). Application of blocker, but not vehicle, elicited pup-directed aggression in sated mice (Fig. 3v), without affecting feeding (Extended Data Fig. 9c). HCN-mediated inhibition of MPOA neurons is therefore sufficient to switch females to pup-directed aggression.

### MPOA neurons encode an aggressive state

These results suggest that quiescent MPOA neurons promote aggression towards pups. To understand how the biophysical changes associated with hunger and estrous state affect neural function *in vivo*, we performed cellular-resolution calcium imaging during pup interactions (Fig. 4a, b). Using a head-mounted miniature microscope, we tracked the activity of individual MPOA neurons before and after food deprivation (Extended Data Fig. 12). Similar to our findings in brain slices, MPOA baseline activity was significantly lower in Agg^+^ than in Agg^−^ mice (Fig. 4c). This difference was present before food deprivation and therefore likely reflects the influence of estrous state (Pre^+^ vs Pre^−^). Pup chemoinvestigation and grooming-associated MPOA responses were weaker in Agg^+^ mice—potentially reflecting their reduced excitability (Fig. 4d and Extended Data Fig. 13b)—, and their absolute tuning, i.e., the magnitude to which they are activated or inhibited, was reduced (Fig. 4e and Extended Data Fig. 13a).

Nonetheless, MPOA neurons still responded during pup aggression, with roughly equal numbers showing activation and inhibition, resulting in a near-zero population average (Fig. 4f, g). Although many MPOA neurons responded during individual pup-directed aggression episodes (Fig. 4f and Extended Data Fig. 14a), their activity showed sustained activity changes—predominantly inhibition (Fig. 4h)—after the onset of aggression and correlated more strongly with this prolonged aggressive state than with specific aggression episodes (Fig. 4h and Extended Data Fig. 14b). Projecting the population activity onto its first two principal components (PCs) revealed a unique state along PC2 in Agg^+^ mice (Fig. 4i). This state was reliably inferred in an unsupervised manner using a Hidden Markov Model (HMM), which detected the majority (94.9 ± 11.3%) of aggression-associated neural activity episodes (Fig. 4j, k). In addition, a linear support vector machine (SVM) trained solely on PC1 and PC2 accurately predicted the animal’s behaviour—in particular aggression— with high accuracy and at a level comparable to an SVM trained on the full neural dataset (Fig. 4l and Extended Data Fig.14c). Notably, the contribution of individual neurons to this aggression state, indicated as their PC2 loading, was correlated with their capacity to predict pup aggression (Extended Data Fig. 14d). Since male mice are spontaneously infanticidal—even when sated—we also examined the MPOA activity patterns associated with pup-directed aggression in males, observing a similar pup aggression-associated state (Extended Data Fig. 15). Thus, in addition to their role in parental behaviour, MPOA neurons encode a distinct state for pup-directed aggression.

What drives the transition of MPOA population dynamics into this aggression state? By modulating neuronal excitability, hunger and estrous state might alter pup representations in the MPOA. We therefore measured changes in neural responses during pup chemoinvestigation before and after food deprivation, using PC distance as a measure of representational similarity. Agg^+^ mice showed greater shifts in pup representations (Fig. 4m) and PC distance was inversely correlated with aggression latency, indicating that larger changes to pup representations lead to faster onset of pup-directed attacks (Fig. 4n). These results suggest that hunger and estrous state promote an aggression state by altering pup representations in the MPOA.

## Discussion

Combining behavioural, circuit-level and cellular approaches, we demonstrate how hypothalamic neurons integrate hunger and estrous state to drive a switch in social behaviour. We identify HCN channels as molecular integrators of these states in MPOA neurons, where baseline channel expression is dynamically set across the estrous cycle. Interestingly, the behavioural switch is a function of 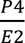 ratio rather than individual hormone levels. Genome-wide targets of Esr1 were recently identified in the brain, including *Hcn1* and the gene encoding PR (*Pgr*)^66^. Intriguingly, E2-administration increases *Hcn1* expression (Extended Data Fig. 11e)^66^, and the chromatin accessibility of *Hcn* genes fluctuates across estrous stages^70^. Although the targets of PR remain less well characterised, it has been shown to inhibit *Esr1*^67–69^. Reciprocal interactions between Esr1 and PR expression, as well as coordinated DNA binding of both receptors^70^, thus likely contribute to hormone ratio sensing. Such sensing may occur in individual MPOA neurons co-expressing PR and Esr1^7^, and/or by neuronal populations with varying hormone sensitivities.

While estrous state modulates *Hcn* expression, food deprivation inhibits HCN channel function via NPY (Fig. 3w). The specific NPY receptor subtypes and downstream signalling pathways remain to be identified, but approximately 70% of MPOA neurons express either NPY receptor Y1 or Y2, both of which inhibit HCN channels by reducing cyclic adenosine monophosphate (cAMP) levels through G_i/o_-protein-coupled mechanisms^71–74^. Knockdown of NPY in Arc^AgRP^→MPOA projections delays, but does not completely abolish, pup-directed aggression (Fig. 3o). This might result from partial AAV transduction efficiency or incomplete knockdown of NPY (Fig. 3i). Along with shorter aggression latencies after prolonged food deprivation (Extended Data Fig. 1a), this delayed aggression phenotype indicates a progressive NPY release during food deprivation. These effects are detectable across the MPOA, including parenting-relevant MPOA^Gal^ neurons (Extended Data Fig. 6).

HCN channels play a well-established role in neuronal excitability^62,74^ and have been found to have a profound impact on states such as sexual satiety and anxiety^75–77^. Reduced HCN function shifts MPOA neurons into a quiescent state with lower baseline activity and excitability (Fig. 3b–d). Aspects of this reduced excitability are also seen *in vivo*, where pup-evoked activity in MPOA neurons is significantly weaker in Agg^+^ mice (Fig. 4c, d). Previous studies have shown that MPOA lesions and optogenetic inhibition induce pup-directed aggression^29,54 37,54,78^, and bulk calcium imaging suggests that MPOA neurons are largely silent during pup attacks in virgin females^78^. These findings promote a view in which aggression primarily results from disinhibition of aggression-promoting neurons downstream of the MPOA^78^. In contrast, our cellular-resolution recordings reveal that most MPOA neurons are either activated or inhibited during aggression, resulting in a minimal net response (Fig. 4f, g). Although MPOA neurons exhibit behavioural tuning—confirming previous work^79^—their activity is even more correlated with a pup aggression state, which can be reliably decoded using both supervised and unsupervised approaches (Fig. 4j, l and Extended Data Fig. 13). Future studies will investigate whether the neurons encoding this state have specific molecular signatures and/or connectivity profiles.

This state-dependent switch may offer behavioural flexibility, enabling adaptive responses to pups during periods of food scarcity, as previously reported for male mongolian gerbils after prolonged food deprivation^80^. Interestingly, food-deprived mice do not appear to perceive pups as food, as pup interactions increase Arc^AgRP^ activity rather than decrease it (Extended Data Fig. 1i–k). This is further supported by the observation that pup and food representations in the MPOA differ after food deprivation (Extended Data Fig. 13l), and that a similar state is present in males during pup-directed aggression (Extended Data Fig. 15). Beyond regulating feeding, Arc^AgRP^ neurons coordinate numerous behavioural adaptations to food deprivation via different projections^5,13,25,30–32,34,39,81–86^. Our findings, along with a recent study^87^, extend their role to the modulation of pup interactions.

A key question in neuroscience and physiology is how internal states interact to drive adaptive behavioural switches^3^. Recent studies have started to address how hunger and thirst jointly regulate ingestive behaviours^88,89^, but how multiple states interact to shape social behaviour remains poorly understood. Our work provides a neural mechanism through which physiological states impart flexibility to pup interactions and offers a conceptual framework for exploring how other states interact to control behaviour.

## Acknowledgments

We thank A. Schaefer, V. Stempel and members of the State-dependent Neural Processing Lab for discussions and comments on the manuscript. We thank the Biological Research Facility at the Francis Crick Institute for animal care and technical assistance, the Crick Light Microscopy, Experimental Histopathology, Mass Spectrometry and Bioinformatics Science Technology Platforms, as well as the Making Lab, Mechanical Workshop, and Vector Core. We thank A. Strosche and V. Stempel for advice on cannulation, H. Fenselau for sharing the hGlyAG construct, S. Wood and A. Resasco for assistance with surgeries, I. Salgarella for assistance with *in vivo* recordings, and N. Borak for sharing implanted mice for miniature microscope recordings. This study received support from the Francis Crick Institute, core funding FC001153. The Francis Crick Institute receives its funding from Cancer Research UK, the UK Medical Research Council, and the Wellcome Trust (J.K.). This study was also supported by European Research Council starting grant ERC-2019-STG847873 (J.K.), NARSAD Young Investigator Award BB/V016946/1 (J.K.), and a Boehringer Ingelheim PhD fellowship (M.C.).

## Author contributions

M.C., R.A. and J.K. conceptualised the study. M.C., R.A., N.L., J.M., M.S. and J.K. contributed to the methodology. M.C., R.A., M.X.C., P.W., A.S., S.L., N.L. and J.K. performed the investigation. M.C., R.A., N.L. and J.K. performed the formal analysis. M.C. and J.K. acquired funding. M.C. and J.K. supervised the project. M.C. and J.K. wrote the original draft of the manuscript. M.C., R.A. and J.K. reviewed and edited the manuscript.

## Methods

### Ethical compliance

All animal procedures performed in this study were approved by the UK government (Home Office) and by the Crick Institutional Animal Welfare Ethical Review Panel (AWERB).

### Mice

Animals were housed in individually ventilated cages on a 12 h:12 h light:dark cycle (light on: 22:00–10:00) with food and water available *ad libitum*. Standard mouse chow (2018S Teklad Global 18% Protein Rodent Diet) was used in all experiments. Baseline (Pre) behavioural testing was performed in the first 4 h of the dark phase and testing after food deprivation (Post) was performed 6 h after the start of the Pre phase, unless stated otherwise. C57BL/6J mice from the Crick breeding colonies were used at age 8–14 weeks for all behavioural experiments. *AgRP-Cre* mice^60^ (JAX #012899) were used to target Arc^AgRP^ neurons. For slice physiology experiments, this line was crossed to Cre-dependent *Rosa26 Tomato* mice (Ai9, JAX #007909). For hormone receptor KO experiments, *Esr1^loxP^* (estrogen receptor α conditional knockout, imported from EMMA, EM:11179)^53^ or *PR^loxP^* (progesterone receptor conditional knockout, made in-house)^7^ were used. All lines were maintained in a C57BL/6J background.

### Behavioural profiling

Virgin females without prior pup exposure were used in all experiments. For experiments in pregnant females, virgin females were paired up with an experienced stud male until a vaginal plug was detected, which marked pregnancy day 1 (D1). Behavioural scoring and analysis were performed by an individual blind to the experimental condition of the animal (e.g., Pre vs Post, manipulation vs control).

#### Pup-directed behaviour assay

Animals were individually housed for 4 days before behavioural testing. Experiments were performed in the home cage and were preceded by a 10 min habituation period. Two C57BL/6 J pups 1–3 days of age were placed in different corners opposite the nest, and pup interactions were recorded for 15 min with a Basler Ace GigE, acA1300-60gmNIR camera. Videos were acquired at a frame rate of 30 Hz using a custom protocol written in Bonsai (Neurogears) and behaviours were scored using Behavioral Observation Research Interactive Software (BORIS)^90^. Pup-directed behaviours were quantified as follows: Contact latency was defined as the time elapsed until the first contact of the test animal’s nose with a pup. Pup grooming was defined as physical contact with pups involving licking, pup displacement and rhythmic head movements. Pup chemoinvestigation was defined as close interaction with the nose of the animal touching the pup but no additional physical contact. The onset of pup retrieval was defined by the time elapsed until a pup was picked up and retrieved to a nest. Time in nest was defined as the time the female stayed in the nest with at least one pup. Crouching was defined as the female stationarily positioned over pups in the nest. Total parenting time is the sum of time spent grooming pups, retrieving pups and time spent in the nest with at least one pup. Nest building was defined as collecting bedding or nesting material and bringing it to the nest as well as shaping it into a new nest. Aggressive contact was defined as close interactions with pups involving rapid, rhythmic head movement, biting, or frantic carrying of pups around the cage. Aggressive animals consistently targeted all pups in the cage. Assays were terminated immediately upon observing pup-directed aggression, and, in rare instances where injury occurred, injured pups were sacrificed. When multiple aggressive contacts are reported, this is due to the introduction of new pups into the cage.

#### Prey assay

House crickets (*Grillus domesticus*, 12–20 mm length, purchased from Northampton Reptile Centre) were used as targets. Immediately after pup-directed behaviour assays, a cricket was placed into the cage for 15 min. Capturing, biting, or biting with forepaw assistance was classified as prey-directed aggression.

#### Residence-intruder assay

Male or female adult mice 8–14-week of age were introduced into the resident’s cage immediately following the pup-directed behaviour assays in randomized order, and resident mice were allowed to interact with the intruder for 15 min. Trials in which the intruder exhibited aggression towards the resident were excluded. Mice were categorised as aggressive towards the intruder if biting and fighting occurred.

#### Elevated plus maze test

A standard elevated plus maze with two closed and two open arms, elevated 90 cm above ground, was used^91^. The assay was initiated by placing the mouse in the open arm of the plus maze, and animal trajectories were recorded for 10 min. Videos were captured and analysed using EthoVision XT 14 (Noldus).

#### Open-field test

A white behaviour test box (60 × 60 × 30 cm, length × width × height) was virtually divided into a centre (30 × 30 cm) and a periphery. A mouse was placed in the periphery, and recorded for 10 min to measure the time spent in the centre or peripheral area. Videos were captured by a top camera and analysed using EthoVision XT 14 (Noldus). Custom detection profiles were set for each mouse, and the detection threshold adjusted so that the mouse could be detected in >95% of video frames. Time spent in the closed vs open arm, and centre vs periphery, average speed, and total time spent moving were quantified using the EthoVision animal tracking pipeline.

#### Food intake

Mice were single housed for 4 days before food intake was measured. On the day of food intake measurement, animals were provided with fresh bedding to avoid leftover food crumbs in the cage. A petri dish with food pellets was provided and 1-h food intake was quantified by calculating the weight difference of the Petri dish. Food intake on behavioural testing days was measured immediately after pup interactions. Baseline food intake was quantified on the day before behavioural testing during the same circadian time in sated mice. Food intake was measured 4 h before the end of the dark phase. For re-feeding, Agg^+^ mice were provided with food *ad libitum* for 1 or 2 h before pup interactions were assessed.

### Mass spectrometry

Trunk blood was collected into EDTA tubes and samples were centrifuged at 2,000 × g for 10 min at 4°C using a microcentrifuge. The supernatant (serum) was pipetted into a fresh 1.5 ml tube and samples stored at −80°C. Ten µl of serum was mixed with 30 μl ice-cold MeOH to induce protein precipitation. Samples were briefly vortexed, put on ice for 5 min and centrifuged at 4°C for 10 min. Thirty µl of extract was mixed with 270 μl MeOH, and 30 µl of the diluted extract were transferred to a vial equipped with an insert, followed by the addition of 1 nmol Scyllo-inositol (Sigma).

Samples were dried and derivatised with 20 μl of freshly prepared methoxyamine (20 mg / ml, in pyridine) (both Sigma) at room temperature for >10 h, followed by a second step of derivatisation with 20 μl BSTFA + 1% TMCS (Sigma) performed at room temperature for 1 h. Data acquisition was performed largely as previously described^92^, using an Agilent 7890B-7000C GC-MSD in EI mode. GC-MS parameters were as follows: carrier gas, helium; flow rate, 0.9 ml / min; column, DB-5MS (Agilent); inlet, 270°C; temperature gradient, 70°C (2 min), ramp to 295°C (12.5°C / min), ramp to 320°C (25°C / min, 3 min hold). Scan range was m/z 50–550. Data analysis was performed using MANIC software version 3.0.20^93^. Metabolite was identified and quantified by comparison to authentic standard of Ghrelin (Anaspec AS-24160).

### Estrous cycle staging

Vaginal smears were taken immediately after pup interaction assays. Animals were scruffed and 20 μl of PBS was gently pipetted several times at the surface of vagina. Samples were air-dried and stained with 10 μl of crystal violet (C.I. 42555, Merck). Mouse IDs were shuffled, and estrous cycle was assessed by an individual blind to aggression phenotype (see ref. ^94^).

### Histology and immunostaining

#### Perfusion and tissue sectioning

Animals were transcardially perfused with phosphate-buffered saline (PBS) followed by 4% paraformaldehyde (PFA) in PBS. Brains were dissected and post-fixed in 4% PFA o/n at 4°C, then washed in PBS. After embedding in 4% low-melting point agarose (Thermo Fisher, 16520-050) in PBS, 60 μm coronal sections were cut on a vibratome (Leica) and mounted on Superfrost Plus slides (VWR, 48311-703) with DAPI-containing Vectashield mounting medium (Vector Laboratories, H-1200). Acute, 250 µm thick, brain sections from electrophysiological recordings were post-fixed in 4% PFA in PBS with 200 mM sucrose (Sigma-Aldrich, S5016) and 0.1 M HEPES (Sigma-Aldrich, H3375) at 4°C on a nutator o/n, rinsed in PBS and washed in PBS-T (0.3% Triton X-100 in PBS) for 1 h.

#### Immunohistochemistry

Immunostaining was performed in 48-well tissue culture plates. Brain sections were permeabilized for 30 min in PBS-T (0.3% Triton X-100 in PBS), post-fixed with 4% PFA in PBS for 10 min and washed in PBS (3 × 20 min). Blocking was carried out for 3 h at room temperature in blocking buffer (3% BSA, 2% normal donkey serum in PBS). Incubation with primary antibodies (in PBS) was performed for 24–48 h on a nutator at 4°C. After washing in PBS (3 × 20 min), secondary antibodies were added in PBS-T for 48 h at 4°C. After final washes in PBS-T (3 × 20 min), sections were mounted. Primary antibodies: rabbit anti c-Fos (Synaptic Systems 226003, 1:2,000), rabbit anti-NPY (Abcam ab30914, 1:500); secondary antibodies: donkey anti-rabbit Alexa Fluor-568 (Thermo Fisher A-11057, 1:2,000), donkey anti-rabbit Alexa Fluor-647 (Thermo Fisher A-21245, 1:2,000).

#### *In situ* hybridisation

Animals were perfused transcardially with ice-cold PBS, and freshly dissected brains embedded in OCT (Tissue-Tek, 4583), frozen on dry ice and stored at −80°C. Subsequently, 18 µm cryosections were cut on a Leica CM1950 cryostat and collected on Superfrost Plus slides (VWR, 48311-703) in three series, only one of which was stained and imaged. Slides were fixed in 10% neutral buffered formalin (NBF), followed by a series of dehydration steps in ethanol (5 min each of 50%, 70%, 100%, 100% v/v EtOH). Slides were pre-treated with RNAscope Protease III reagent for 30 min at 40°C. Single-molecule fluorescent *in situ* hybridisation were performed on slides using the RNAscope LS Multiplex Reagent Kit (Advanced Cell Diagnostics), LS 4-Plex Ancillary Kit and Multiplex Reagent Kit on a robotic staining system (Leica BOND-III). RNAscope probes were Hcn1 (ACD, Cat. No. 423658), Hcn2 (Cat. No. 427009), Hcn3 (Cat. No. 551528) and Hcn4 (Cat. No. 421278). Immunostainings against the neuronal marker NeuN were subsequently performed (Millipore MAB377, 1:500).

#### Imaging

Images were acquired on a Vectra Polaris Automated Quantitative Pathology Imaging System (Akoya Biosciences) at 20× magnification. Regions of interest (ROIs) were selected using Phenochart software (Akoya Biosciences) and image tiles were spectrally unmixed using inForm Tissue Analysis software (Akoya Biosciences). Stitching of spectrally unmixed image tiles and image analyis were performed in QuPath software^95^. C-Fos-positive nuclei (or NeuN-positive neuronal cell bodies) were first detected using custom QuPath scripts. Detection of Esr1, PR and HCN transcripts was subsequently performed on cell body detections. Thick brain sections (250 µm) were imaged on an upright confocal microscope (Zeiss LSM 710) using a 63× (NA 1.4) oil immersion objective and a Z step size of 0.5 μm.

#### Constructs for NPY knockdown

Constructs for shRNA-mediated knockdown of *Npy* were developed using the Broad Institute’s hairpin design tool (https://portals.broadinstitute.org/gpp/public/seq/search) on the *Npy* coding sequence (NM_023456.3, position: 3728–3748). The following sequences were used: (1) Npy_817 gaagttataagccttgtttCACTGATTTCAGACCTCTTAA CTCGAG TTAAGAGGTCTGAAATCAGTG TTTTTactagttactaggatccg; (2) Npy_818 gaagttataagccttgtttGCTCTGCGACACTACATCAAT CTCGAG ATTGATGTAGTGTCGCAGAGC TTTTTactagttactaggatccg (uppercase = stem-loop-stem; TTTTT = termination signal). Both were cloned into pAAV-G-Creon shRNA[Control] (Addgene #181824)^96^ which yielded the constructs pAAV-G-CreON-shRNA_817-NPY-GFP and pAAV-G-CreON-shRNA_818-NPY-GFP, respectively. As a negative control, a scrambled sequence (CCTAAGGTTAAGTCGCCCTCGctcgagCGAGGGCGACTTAACCTTAGGTTTTTT) was designed using VectorBuilder.

### Surgical procedures

Analgesia was provided one day before surgery (0.15 ml carprofen in 200 ml drinking water). Mice were anaesthetized using isoflurane (4% for induction, 1.5% for maintenance) in oxygen-enriched air and head-fixed in a stereotactic frame (Model 940, Kopf Instruments). Meloxicam (10 mg / kg body weight) and Buprenorphine (0.1 mg / kg body weight) were given subcutaneously before craniotomy. The surgery site was closed using Vicryl sutures (Ethicon) or Vetbond surgical glue (3M). Carprofen was provided in drinking water for 2 days following surgery for postoperative pain management. Eyes were protected with ophthalmic ointment (Viscotears, Alcon). Rectal body temperature was maintained at 37°C during surgery using a heating pad (Harvard Apparatus) and animals were kept in a heated recovery chamber until fully mobile. Animals were allowed to recover for at least 2 weeks before behavioural testing.

#### Brain coordinates

See Extended Data Table 1 for injection, implantation and recording coordinates. Chemogenetic effectors were injected into two rostrocaudal Arc coordinates (–1.4 / ± 0.25 / –5.90 and –1.6 / ± 0.25 / –5.90 mm) to maximise the number of transduced neurons. For projection-specific *Npy* knockdown, MPOA coordinates were adjusted to 0.0 / ± 0.3 / –5.05 mm to maximise the number of retrogradely labelled Arc^AgRP^ neurons.

#### Chemogenetics

For chemogenetic activation, 200 nl of AAV5-hSyn-DIO-hM3Dq(Gq)-mCherry (Addgene #44361, 2.5 × 10^13^ GC / ml) or AAV5-hSyn-DIO-mCherry (Addgene #50459, 1.8 × 10^13^ GC / ml) was injected into the Arc (see Extended Data Table 1 for coordinates). After assessing animals’ spontaneous pup-directed behaviours 3 weeks after viral injection, CNO (Bio-Techne 12352200, 3 mg / kg) was injected intraperitoneally, and pup-directed behaviour assessed 30 min later. For chemogenetic inhibition, 250 nl of AAV5-loxP-hGlyAG-2A-nlsVenus (1.6 × 10^13^ GC / ml, Crick Vector Core) was prepared from a pAAV-loxP-hGlyAG-2A-nlsVenus plasmid^20,21^ (gift from H. Fenselau) and injected into the Arc (see ‘Brain coordinates’). Ivermectin (5 mg / kg, dissolved in 7 : 3 propylene glycol : glycerol) was injected 24 h before the start of food deprivation, and behaviour assessed after 6 h of food deprivation.

#### Optogenetics

To optogenetically activate Arc^AgRP^ projections, 250 nl of AAV5-EF1a-DIO-ChR2-EYFP or AAV1-EF1a-DIO-ChR2-EYFP (Addgene #20298, 0.7 × 10^13^ GC / ml) or AAV1-EF1a-DIO-YFP (Addgene #27056, 2.5 × 10^13^ GC / ml) were injected into the Arc (see Extended Data Table 1 for coordinates). During the same surgery optic fibers (Doric Lenses) were implanted 200–400 µm above the target 2area (MPOA: dual fiber cannula 200/245 μm, 0.37 NA, GS1.0; LPOA: dual fiber cannula 200/245 μm, 0.37 NA, GS2.0; PVH: mono fiber cannula 400/470 μm, 0.37 NA). After 2–3 weeks of recovery, animals were connected to matching patch cords connected to a laser (Stradus 473-80 nm, Vortran) via a commutator (RJ1, Thorlabs). Four distinct protocols for optogenetic stimulation were used: acute stimulation whenever animals were close to a pup, or 15 / 30 / 60 min of pre-stimulation (pre) followed by a 15-min pup-directed behaviour assay. A period of 3–4 days was allowed between two consecutive optogenetic experiments to prevent sensitization to pups. The light power exiting the fiber tip corresponded to an irradiance of 4.68 mW × mm^−2^ at the target region (see http://www.stanford.edu/group/dlab/cgi-bin/graph/chart.php). For acute stimulation, blue light was delivered in 20-ms pulses at 20 Hz for 1–4 s whenever the animal contacted a pup with its snout. In the pre stimulation protocols, cycles of 1 s of 20 Hz stimulation followed by 4 s without stimulation were delivered for the indicated duration^23^.

#### Hormone receptor knockout

AAV2/1-syn-fDIO-EGFP-2A-iCre (250 nl) or AAV2/1-syn-fDIO-GCaMP7s-2A-iCre (250 nl) was injected into the MPOA (coordinates as above; custom plasmids, packaged by Vectorbuilder) of *Esr1^loxP^* or *PR^loxP^* mice. Viral titers were adjusted to 2.1 × 10^13^ GC / ml. Animals were tested 3 weeks after injection and brain slices were subsequently prepared for histological analysis. The efficiency of viral-genetic receptor KO was established in an separate experimental cohort of *Esr1^loxP^* or *PR^loxP^* animals which received unilateral MPOA injections of either AAV2/5-CMV-EGFP-Cre (250 nl, Addgene 105545, 2 × 10^13^ GC / ml) or AAV2/5-CMV-EGFP (250 nl, Addgene 105530, 2 × 1013 GC / ml), and which has since been published^7^.

#### NPY knockdown

pAAV-G-CreON-shRNA_817-NPY-GFP and pAAV-G-CreON-shRNA_818-NPY-GFP (see ‘Constructs for NPY knockdown’) were packaged as rAAV2-retro capsids and the titer measured via qPCR. For projection-specific knockdown of NPY, 400 nl of a 1:1 mix of AAV-retro-G-CreON-shRNA_817-NPY-GFP (3.8 × 10^13^ GC / ml) and AAV-retro-G-CreON-shRNA_818-NPY-GFP (2.3 × 10^13^ GC / ml) was bilaterally injected into the MPOA (see ‘Brain coordinates’). As control, AAV-retro-CreON-shRNA-scr expressing a scrambled shRNA (400nl, 1.78 × 10^13^ GC / ml) was injected. Behavioural testing and/or electrophysiological recordings were performed 3 weeks after injection.

#### Cannulation experiments

Mice were implanted with stainless steel bilateral guide cannulas (C235GS-5-1.0/SPC, Protech International) 0.2 mm above the MPOA. Cannulas were fixed to the skull with dental cement. Dummy cannulas (C235DCS-5/SPC, Protech International) were inserted into guide cannulas to prevent clogging and closed with a dust cap. Mice were allowed to recover for 4 days. One hour before behavioural testing (see ‘Pup-directed behaviour assay’), 1 µl of ZD-7288 (Tocris #1000; 1 mM, in sterile artificial cerebrospinal fluid, ACSF) or ACSF alone (vehicle, ctrl) were administered to each side of the cannula at a rate of 0.5 µl / min.

#### Fiber photometry

AAV-hsyn-DIO-GCaMP7s (Addgene 104491-AAV1, 300 nl, 1.5 × 10^13^ GC / ml) was injected into the Arc of *AgRP-Cre* mice and a 200 µm fiber-optic cannula (MFC_200/230-0.37_6mm_MF1.25_FLT, Doric Lenses) implanted into the MPOA (see Extended Data Table 1 for coordinates). The cannula was fixed to the skull using UV light-curable glue (RelyX Unicem, 3M) and Superbond cement (Prestige Dental). Recordings were performed 3 weeks after surgery using an FP3001 fiber photometry system (Neurophotometrics). Briefly, two LEDs (415 nm, 470 nm, light power ∼50 µW) were pulsed at 20 Hz in an interleaved manner to obtain an isosbestic motion signal (415 nm) and GCaMP activity (470 nm). A FLIR 277 BlackFly CMOS camera was used to detect fluorescent signals, and acquisition was controlled (and synchronised to the acquisition of behavioural video recordings) via Bonsai (https://bonsai-rx.org/).

#### Miniature microscope imaging

AAV2/1-syn-GCaMP7s (Addgene 104487, 100–200 nl, 2 × 10^13^ GC / ml) was unilaterally injected into the MPOA of C57BL/6J mice using a Nanoject II or Nanoject III injector (Drummond Scientific) and pulled glass capillaries (3-000-203-G/X, Drummond Scientific). See Extended Data Table 1 for injection and implantation coordinates. After letting the virus diffuse for 5 min, the injection needle was slowly retracted and an integrated gradient-index (GRIN) lens (0.6 × 7.3 mm, 1050-002177, Inscopix) was slowly implanted and fixed to the skull using UV light-curable glue (RelyX Unicem, 3M) and Superbond cement (Prestige Dental). Recordings started 6–8 weeks after surgery. Mice were connected to a miniature microscope (nVista, Inscopix) to check for sufficient expression of GCaMP7s. Imaging data were acquired using nVista HD software (Inscopix) at a frame rate of 20 Hz with 475 nm LED power of 0.1–0.2 mW / mm^2^, analog gain of 5–8, and an image resolution of 800 × 1,280 pixels. Imaging parameters and focal depth were kept identical across sessions. Imaging and behavioural video collection was synchronized using Bonsai software (Neurogears). Mice were connected to the microscope and allowed 20 min of habituation before recordings were performed in their home cage. A 1-min baseline was acquired before introducing pups, which was used to calculate the relative fluorescence change for each ROI in the field of view.

### *Ex vivo* electrophysiology

C57BL/6J mice were deeply anaesthetized with 3% isoflurane in oxygen and decapitated. The brain was quickly dissected and placed in ice-cold slicing solution containing (in mM): sucrose (213), KCl (2.5), NaH_2_PO_4_ (1.3), NaHCO_3_ (26), MgSO_4_ (2), CaCl_2_ (2), and D-glucose (10), equilibrated with carbogen (95% O_2_ / 5% CO_2_). Coronal brain slices (250 μm thickness) containing the MPOA were cut on a vibratome (Leica VT1200S) in ice-cold slicing solution and transferred to an incubation chamber with artificial cerebrospinal fluid (ACSF) containing (in mM): NaCl (127), KCl (2), NaH_2_PO4 (1.2), NaHCO_3_ (26), MgSO_4_ (1.3), CaCl_2_ (2.4) and D-glucose (10), which was continuously oxygenated with carbogen. After at least 1 h of recovery at 37°C, slices were transferred to a submersion chamber under an upright microscope with infrared Nomarski differential interference contrast optics (Slicescope, Scientifica). During recordings, slices were submerged in, and continuously perfused (2–3 ml / min) with, ACSF at near physiological temperature (33°C) and continuously oxygenated with carbogen. Glass micropipettes (3–6 MΩ resistance) were pulled from borosilicate capillaries (World Precision Instruments, Aston, UK) on a P-97 Flaming/Brown micropipette puller (Sutter, Novato, CA) and filled with internal solution containing (in mM): K-Gluconate (140), KCl (10), KOH (1), EGTA (1), Na_2_ATP (2), HEPES (10), pH 7.3, 280–290 mOsm. Access resistance was monitored throughout the experiment, and neurons in which it exceeded 25 MΩ or changed by ≥ 20% were excluded. Liquid junction potential was 16.4 mV and not compensated. We characterized the intrinsic electrophysiological properties of cells using a standardized current-clamp protocol that consists of I/V curves, ramps, and current injections. HCN-mediated voltage sag amplitude was measured in response to hyperpolarising 1-s direct-current steps^97^. T-Type calcium currents (CaT) were assessed by a standard current clamp protocol in which cells were hyperpolarised to −120 mV and then stepped back to −60 mV^98^. The amplitude of the resulting rebound was then quantified. To assess excitability, ramping depolarizing currents (10 pA / s) from +25 to +165 pA were injected. Spontaneous postsynaptic currents (sPSCs) were detected using a threshold-based detector (WinEDR v4, template mode). Rise time is defined as the time needed for sPSC amplitude to reach *1-e*^-1^ (≈ 63%) of its maximal value and time constant of decay defined as the time needed for sPSC amplitude to return to *1/e* (≈ 37%) of resting state. The HCN channel blocker ZD-7288 (Tocris #1000) was added at a concentration of 50 µM 1 h before the start of recordings. NPY (Phoenix Pharmaceuticals 049-03) was added at a concentration of 100 µM 1 h before the start of recordings. NPY receptor antagonists (NPY1R: 10 µM BIBP 3226, Tocris 2707; NPY2R: 100 nM BIIE 0246, Merck SML2450) were added to brain slices 1 h before the start of recordings^99,100^. Recordings were acquired using a Multiclamp 700B amplifier (Molecular Devices), low pass filtered at 10 kHz and digitized using a Digidata 1322A digitizer (Molecular Devices). Slow and fast capacitive components were semi-automatically compensated. Offline data analysis was performed with Clampfit 10 software (Molecular Devices), WinEDR v4, WinWCP v5 (http://spider.science.strath.ac.uk/sipbs/software_ses.htm), and custom routines written in Python 3.7.

#### Channelrhodopsin-assisted connectivity mapping

For channelrhodopsin-assisted connectivity mapping^101^, 200 nl of AAV1-EF1a-FLEx-hChR2(H134R)-EYFP (Addgene 20296, 7 × 10^12^ GC / ml) was bilaterally injected into the MPOA of *AgRP-Cre* mice. Acute brain sections were prepared 3 weeks after viral injection. We used a CsCl based internal solution containing (in mM): CsCl (140) Hepes (10), EGTA (1), Na_2_ATP (2), HEPES (10), pH 7.3, 280–290 mOsm. Spontaneous inhibitory postsynaptic currents (sIPSCs) were recorded in voltage clamp configuration at –70 mV in presence of 1 µM TTX (Alomone T-550) and 100 μM 4-AP (Sigma 275875). Drugs were washed in at least 10 min before starting recordings. Photostimulation was delivered from a 490 nm LED (pE-100, CoolLED) through a 60× objective and consisted of 2–10 ms light pulses, at a light intensity of ∼2.6 mW / mm^2^.

### Quantification and data analysis

Unless stated otherwise, data are presented as mean ± s.e.m. Box plots represent the median and 25^th^ and 75^th^ percentiles, and whiskers represent the data range. Significance levels are: \*\*\**P* < 0.001; \*\**P* < 0.01; \**P* < 0.05.

#### Calculation of predicted baseline switching rates

The observed switching rates of animals with hormone receptor ablation were compared to the predicted baseline switching rate which would be expected for each cohort if receptors were intact. These baseline rates were determined using hypothesis testing on Poisson binomial distributions, which were constructed based on the estrous cycle distribution of each cohort. The mode of each distribution constituted the expected switching rate which was to the observed switching rates.

#### Image analysis and registration

The Fiji plugin ABBA (https://biop.github.io/ijp-imagetoatlas/) was used to register coronal brain sections to the Allen Brain Atlas (CCFv3)^102^. Briefly, X and Y rotation were adjusted across all sections from a given brain and two rounds of affine registration using Elastix were performed. Samples then underwent non-rigid registration using the BigWarp tool (sample channel: DAPI, atlas channel: Nissl). Positive cell detection was performed on the transformed samples in QuPath. Transformed cell detections were exported from QuPath, visualized using a custom Python app (https://github.com/nickdelgrosso/ABBA-QuPath-RegistrationAnalysis), and analysed using custom Python scripts.

#### Quantification of NPY knockdown efficiency

Brain sections from *AgRP-Cre* mice injected with conditional AAVs expressing GFP and NPY-targeting shRNA (or negative controls, see ‘NPY knockdown’) were immunostained against NPY (see ‘Immunohistochemistry’) and imaged on a confocal microscope (Zeiss LSM 710). Pixel-based analysis was performed because NPY staining is not exclusively localised to neuronal cell bodies. Image stacks were imported into ImageJ, and the JaCoP plugin^103^ was used to calculate the percentage of pixels in the NPY channel that co-localised with GFP-positive pixels.

#### Processing and analysis of fiber photometry data

The recorded interleaved trace was split into isosbestic (415 nm) and activity (470 nm) signals using custom Python routines. A linear fit of the isosbestic signal was calculated and subtracted from the activity channel to remove baseline fluorescence and motion artifacts. An additional moving minimum baseline (20 s sliding window) was subtracted from each resulting trace to account for slow fluctuations in activity, such as bleaching. The relative fluorescence change was calculated as 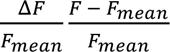 and was normalized using a min max scaler. Manually scored behaviours were aligned to the activity traces via timestamps acquired in Bonsai (Neurogears).

#### Processing and analysis of miniature microscope imaging data

##### Pre-processing

Image frames were spatially downsampled to 400 × 540 pixels. Drift of the baseline signal over time was removed using a spatial bandpass filter with lower and upper cut-off spatial frequencies of 0.005 and 0.5 oscillations per pixel, respectively. Motion artifact correction was performed, and the relative fluorescence change Δ*F*/*F*_0_ for each pixel compared to baseline calculated as 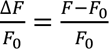 (where *F*_0_ is the mean fluorescence value of each pixel during the baseline period). PCA/ICA-based cell detection was performed using a mean ROI radius of 7–9 pixels in the Inscopix Data Processing Software. All automatically identified cells were manually verified to exclude false-positive detections, and cells not detected by the algorithm were manually added. Cell traces were deconvolved using OASIS^104^ with model order 1 and a spike SNR threshold of 3.0.

##### Longitudinal registration

Longitudinal registration of Pre and Post field-of-views was performed using Inscopix Data Processing software without session correlation for thresholding. The resulting aligned traces were manually quality-controlled. Identified ROIs with non-round shapes or without any visible evoked activity were discarded

##### Evoked activity analysis

*Z* scores were calculated based on the period of –2 to 0 s before event detection (baseline period) and 0 to 2 s from event detection (activity period) as *z* = *x* − *µσ*, where *x* is Δ*F*/*F* of the current timestamp, *µ* is the mean Δ*F*/*F* of the baseline period and *σ* is the standard deviation of the baseline period. Significant responses were called when the *Z*-scored Δ*F*/*F* of the baseline and activity periods were significantly different (unpaired *t* test). ROIs were thereafter categorized as exhibiting increased, decreased, or unchanged evoked activity. The single neuron tuning index was derived from performing an unpaired *t* test between the activity and baseline periods. Tuning indices were normalized based on their absolute value using a min max scaler from the *sklearn* preprocessing package while keeping their original sign.

##### PCA and classifier training

For the calculation of principal component (PC) distances, episodes of 5 s after the onset of indicated behaviours were extracted from each neuron. All episodes were standardized prior to analysis. Principal components cumulatively accounting for 90% of the variance in the episodes were utilized for these calculations. For training the PC-based SVM classifier, all behavioural episodes of >2 s were selected. The training and testing samples were split by episode number rather than by frame number.

## Extended Data Figures

**Extended Data Fig. 1:**
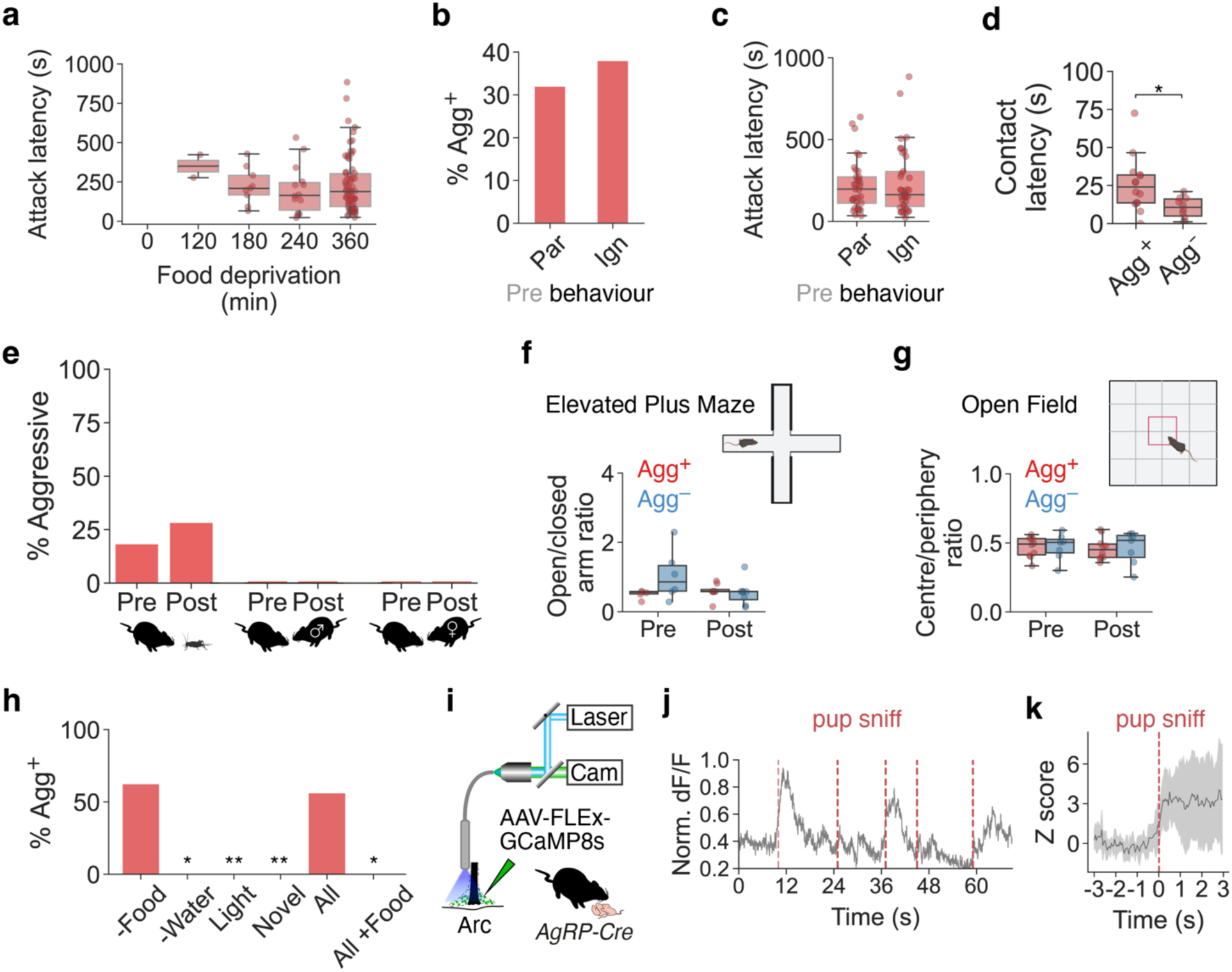
State- and target-specificity of the negative parental switch. **a**, Attack latency as a function of food restriction duration (n = 4, 9, 14, 70 mice, *P* = 6.5 × 10^−20^). **b**, Percentage of Agg^+^ animals depending on behaviour before food deprivation (Par, parental, Ign, ignoring; n = 67, 66 mice). **c**, Attack latency depending on behaviour before food deprivation (n = 33, 37 mice). **d**, Pup contact latency of Agg^+^ and Agg^−^ mice (n = 33, 37 mice). **e**, Percentage of Agg^+^ mice exhibiting aggression towards prey (cricket) or adult intruders in Pre and Post period (n = 16, 10, 10). **f**, Performance of Agg^+^ and Agg− mice in Elevated Plus Maze before and after food deprivation (n = 6, 8). **g**, Performance of Agg^+^ and Agg^−^ mice in Open Field before and after food deprivation (n = 7, 10). **h**, Percentage of mice switching to pup-directed aggression after 6 h of either food restriction, water restriction, light cycle inversion, or housing in novel, empty cage. All, all stressors, All+Food, all stressors in non-food-deprived mice (n = 16, 8, 11, 10, 16, 8). Significance levels are between ‘–Food’ and all other groups. **i**, Fiber photometry recordings from Arc^AgRP^ neurons during pup interactions. **j**, Example recording trace of Arc^AgRP^ population activity during pup chemoinvestigation in 6-h food-deprived mice. **k**, Average, *Z*-scored Arc^AgRP^ activity during pup chemoinvestigation (5 traces from n = 1 mouse). One-way ANOVA in **a**. Fisher’s exact test in **b, e, h**, with Benjamini-Hochberg adjustment in **h**. *U* test in **c, d**. Two-way ANOVA in **f, g**.

**Extended Data Fig. 2:**
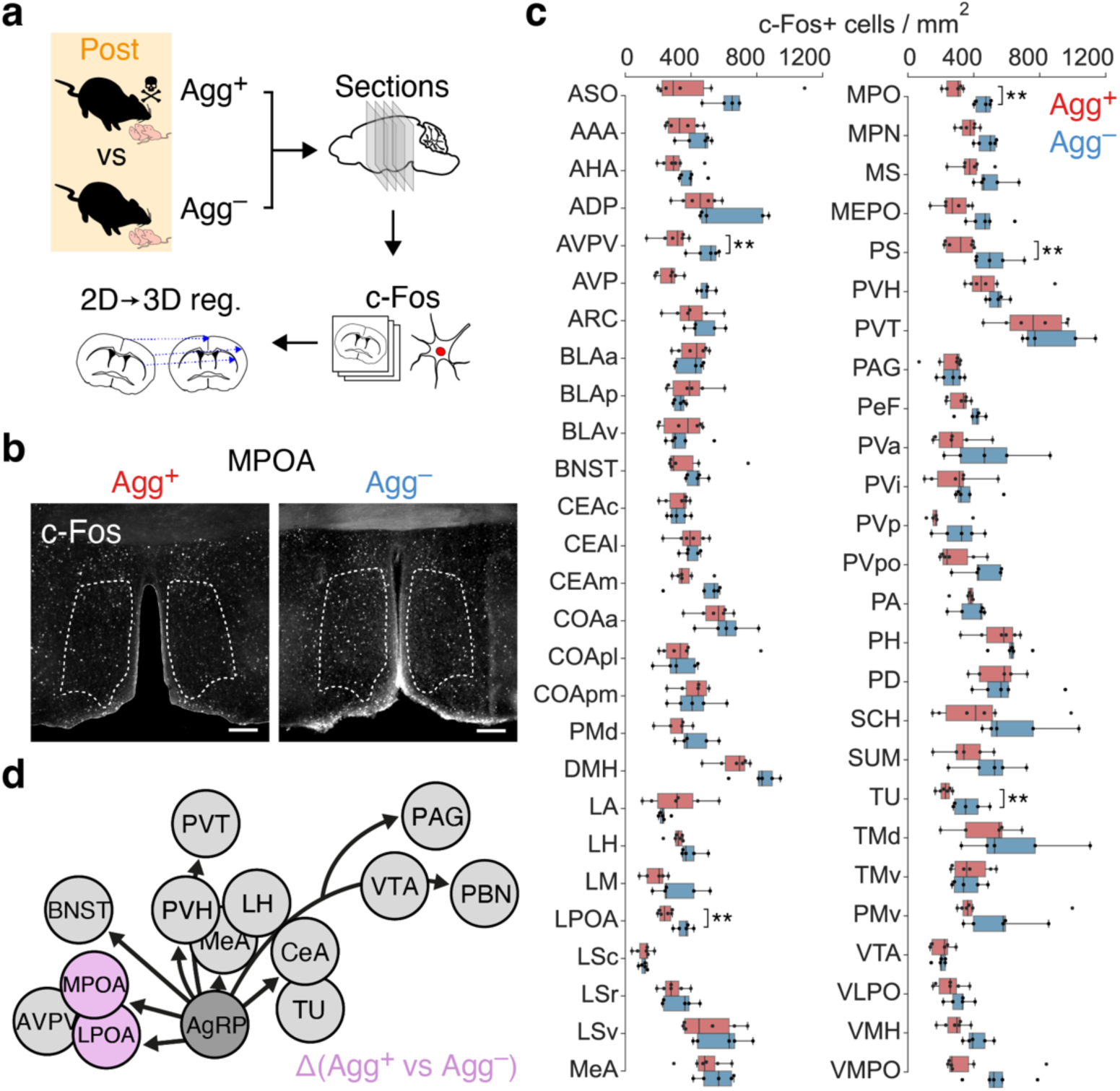
Identification of candidate Arc^AgRP^ targets via activity mapping. **a**, Immunostainings against c-Fos in brain sections from Agg^+^ and Agg^−^ mice, and registration to Allen Brain Atlas (see Methods). **b**, Example c-Fos^+^ cell densities in MPOA of Agg^+^ and Agg^−^ mice in MPOA sections. Scale bars, 200 µm. **c**, Density of c-Fos^+^ cells in hypothalamic brain areas of Agg^+^ and Agg^−^ mice (n = 6, 5 mice). **d**, Arc^AgRP^ projections. Areas with significantly different c-Fos^+^ cell numbers between Agg^+^ and Agg^−^ groups, and receiving direct Arc^AgRP^ projections, are highlighted. Note that ADP, MPN, MPO, PD and PS are MPOA subregions. *U* tests with Benjamini-Hochberg adjustment in **c**. See Extended Data Table 1 for acronyms of brain areas.

**Extended Data Fig. 3:**
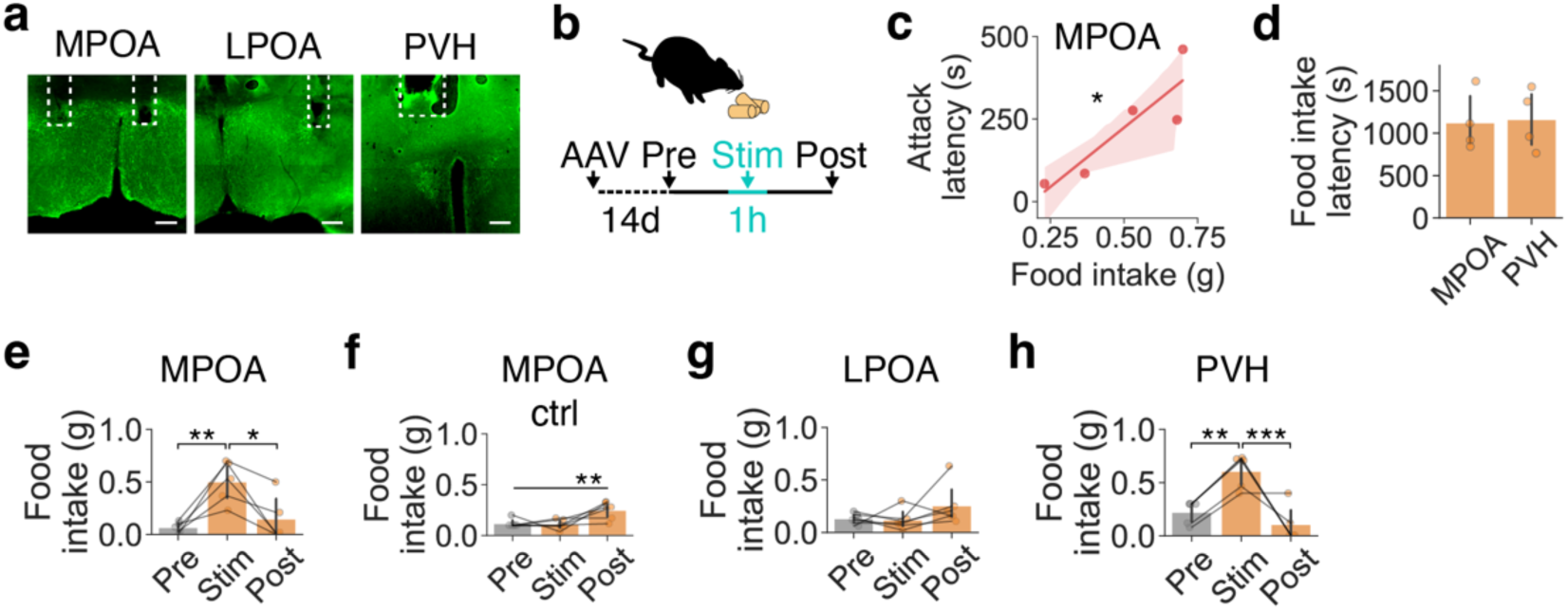
Effects of optogenetically activating Arc^AgRP^ projections. a, Implantation sites of optical fibers in MPOA, LPOA and PVH of *AgRP-Cre* mice injected with AAV-DIO-ChR2-EYFP. Scale bars, 200 µm. b, Optogenetic stimulation paradigm to assess food intake. c, Correlation between food intake and pup attack latency in mice with optogenetic stimulation of Arc^AgRP^→MPOA projections (linear regression, *r*^2^ = 0.784; *P* = 0.046, n = 5 mice). d, Latency of food intake during continuous stimulation of Arc^AgRP^→MPOA and Arc^AgRP^→PVH projections (n = 4, 4 mice). e, f, Effect of optogenetically activating Arc^AgRP^→MPOA projections (e, 1-h pre-stim, n = 5) on 1-h food consumption in sated mice, and negative control (f, *AgRP-Cre* mice injected with AAV-DIO-EYFP, n = 6). g, h, Effect of optogenetically activating Arc^AgRP^→LPOA (g, n = 6) or Arc^AgRP^→PVH (h, n = 5) projections on 1-h food consumption in sated mice. *U* test in d. One-way ANOVA with Tukey *post hoc* test in e–h.

**Extended Data Fig. 4:**
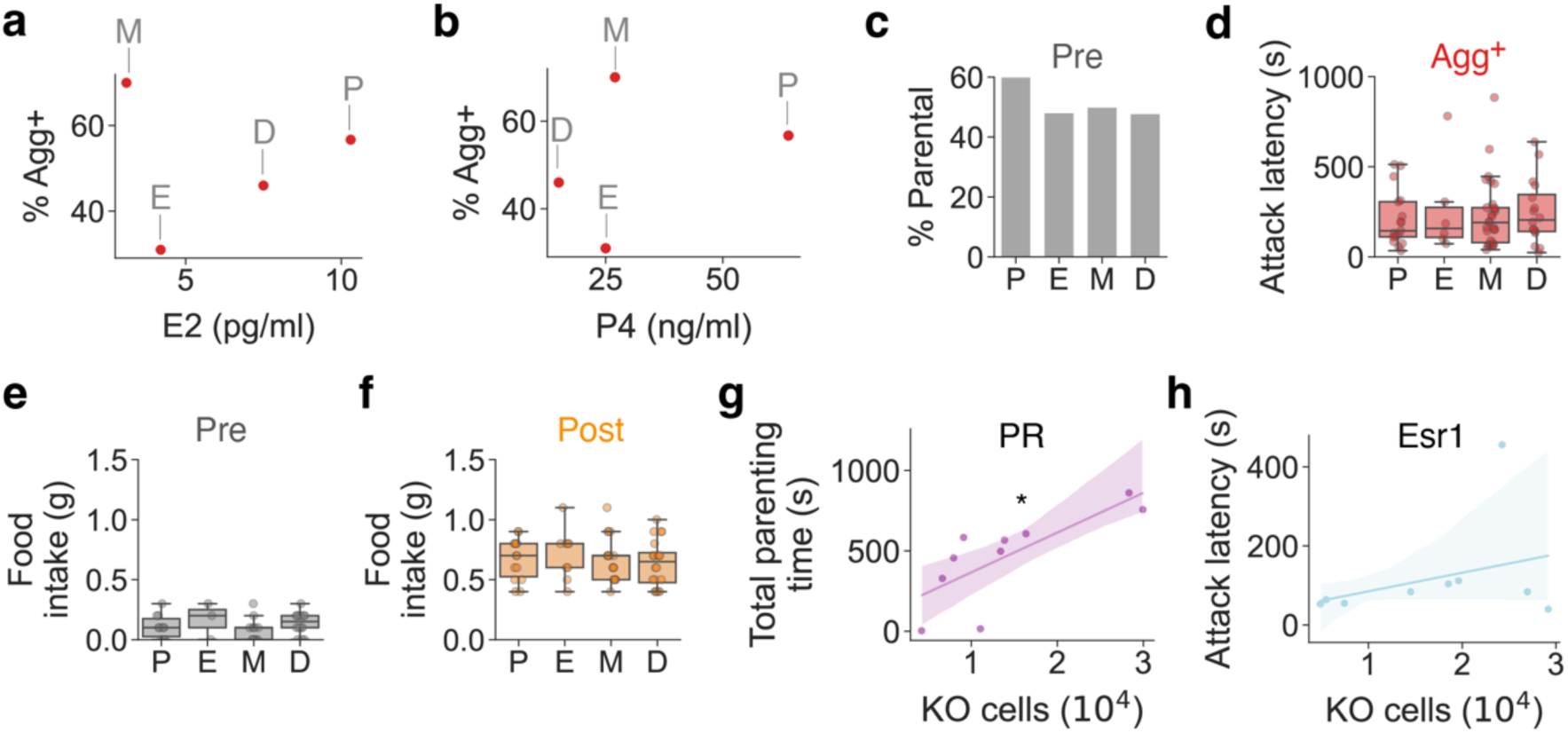
Effects of estrous state on parental interactions and food intake. **a, b**, Switching rate does not correlate with P4 (**a**) or E2 (**b**) plasma concentration (P, E, M, D; n = 30, 19, 40, 37 mice). **c**, Percentage of spontaneously parental virgin female mice in different estrous states before food deprivation (n = 35, 26, 52, 45 mice). **d**, Attack latency of Agg^+^ mice in different estrous states (n = 30, 19, 40, 37 mice). **e, f**, 1-h food consumption in sated mice before (**e**) and after (**f**) food deprivation depending on estrous state (Pre: n = 10, 3, 11, 14; Post: n = 14, 9, 17, 20). **g**, Correlation between total parenting time (see Methods) and number of PR-ablated MPOA neurons (linear regression, *r*^2^ = 0.581; *P* = 0.01, n = 10). **h**, Correlation between attack latency and number of Esr1-ablated MPOA neurons (linear regression, *r*^2^ = 0.113; *P* = 0.377, n = 9). P, proestrus, E, estrus, M, metestrus, D, diestrus. Fisher’s exact test with Benjamini-Hochberg adjustment in **c.** One-way ANOVA in **d–f**.

**Extended data Fig. 5:**
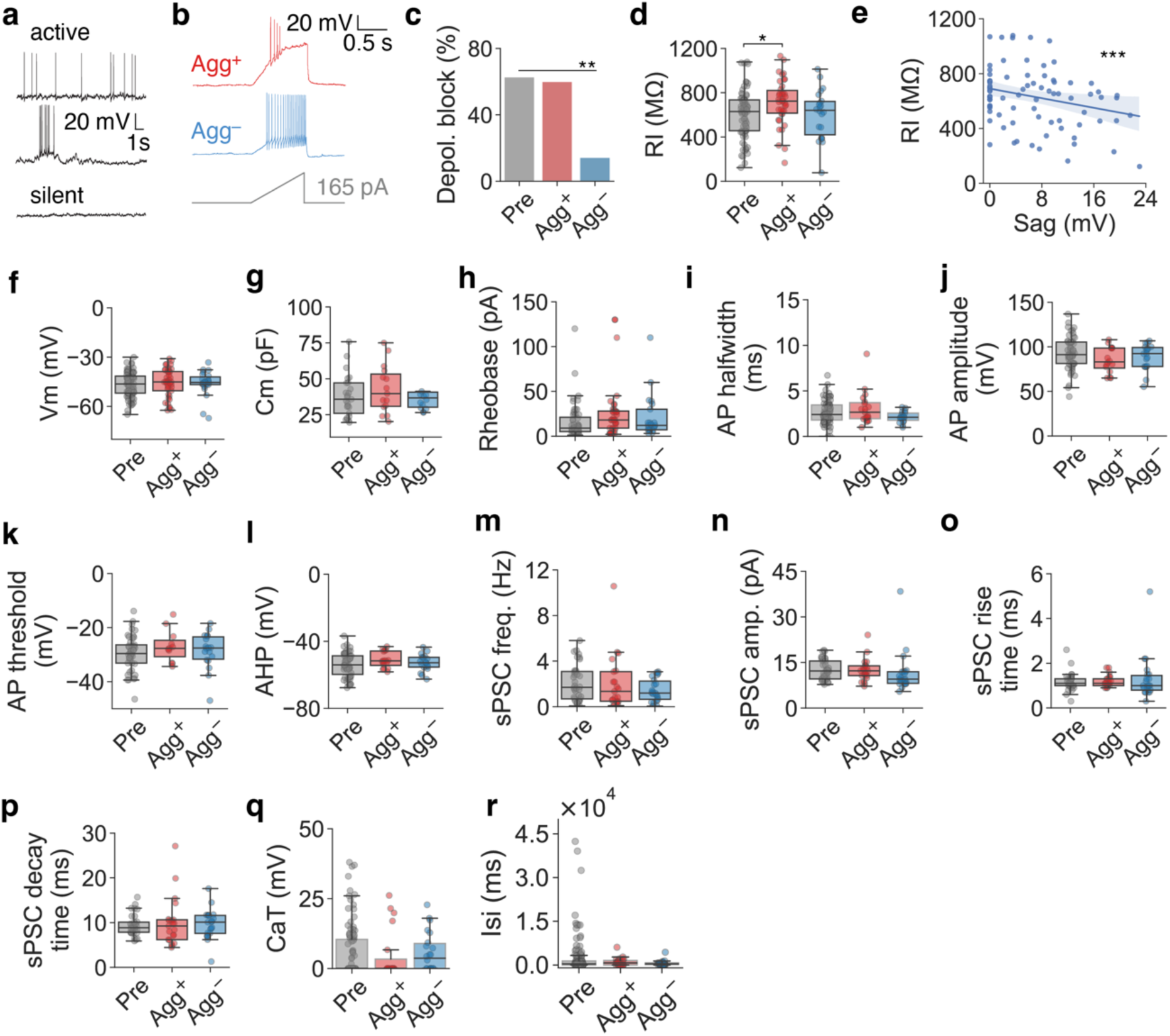
Biophysical effects of food deprivation on MPOA neurons. **a**, Example current-clamp traces of different activity patterns of MPOA neurons at resting membrane potential. **b**, Example current clamp recording traces of cells with (Agg^+^) and without (Agg^−^) depolarization block. **c, d, f–r**, Whole-cell recordings from MPOA neurons in mice before (Pre) and after (Agg^+^ and Agg^−^) 6-h food deprivation (242, 29, 22 cells from n = 24, 7, 3 mice in **c, d, f–i, m–p, r**; 45, 11, 18 cells from n = 24, 7, 3 mice in **j–l, q**): percentage of neurons exhibiting depolarization block (**c**), input resistance (**d**), resting membrane potential (**f**), membrane capacitance (**g**), rheobase (**h**), action potential half-width (**i**), action potential amplitude (**j**), action potential threshold (**k**), afterhyperpolarization (**l**), sPSC frequency (**m**), sPSC amplitude (**n**), sPSC rise time (**o**), sPSC decay time (**p**), rebound after hyperpolarisation due to T-type calcium currents (CaT) (**q**), and inter-spike interval (**r**). **e**, Correlation between input resistance and voltage sag amplitude (linear regression, *r*^2^ = 0.07; *P* = 0.003, n = 74). Chi-Square test with Benjamini-Hochberg adjustment in **c**. One-way ANOVA with Tukey *post hoc* test in **d, f–r**.

**Extended data Fig. 6:**
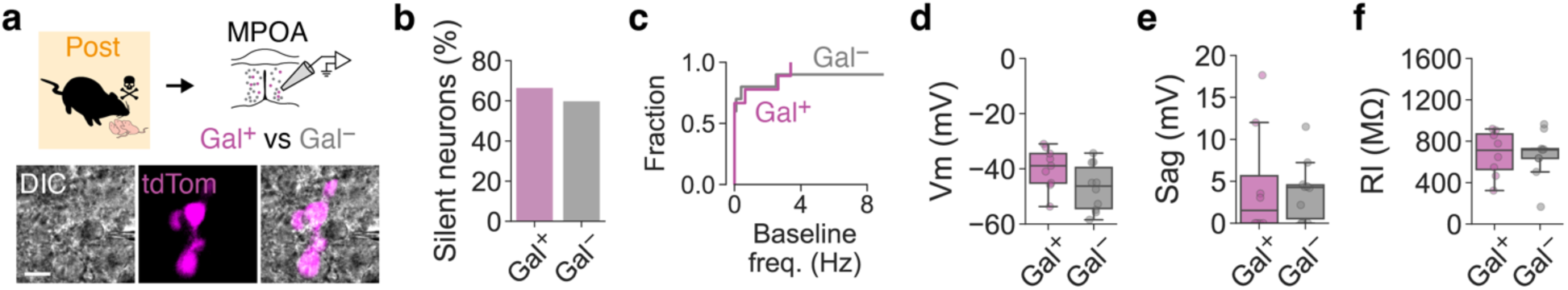
Food deprivation-induced biophysical changes in MPOA^Gal^ neurons. **a**, Whole-cell recordings from Galanin-positive (Gal^+^) and -negative (Gal^−^) MPOA neurons in Agg^+^ mice (11, 10 cells from n = 3, 4 mice). Scale bar, 20 µm. **b–f**, Percentage of silent neurons at resting membrane potential (**b**), baseline firing frequency (**c**), resting membrane potential (**d**), voltage sag amplitude (**e**) and input resistance (**f**) of Gal^+^ and Gal^−^ MPOA neurons. Fisher’s exact test in **b**, *U* test in **c–f**.

**Extended Data Fig. 7:**
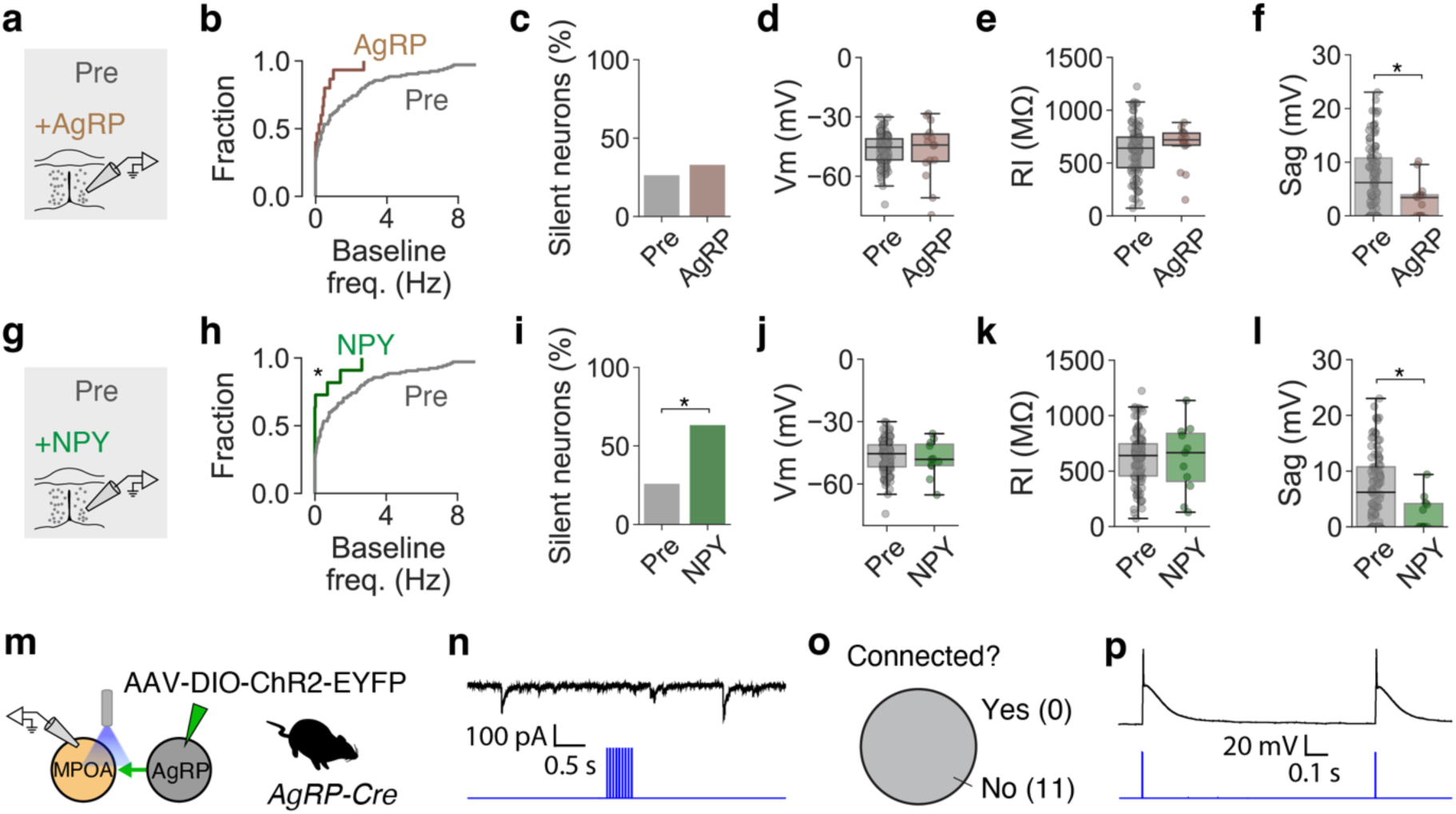
NPY mimics the biophysical effects of food deprivation on MPOA neurons. **a, g**, Recordings from MPOA neurons without (Pre) and with bath application of 100 µM AgRP (**a**, 102, 15 neurons from n = 12, 5 mice) or 100 µM NPY (**g**, 102, 11 cells from 12, 4 mice). **b**–**f** and **g–l**, biophysical parameters of MPOA neurons at baseline (Pre) and with addition of AgRP or NPY: baseline frequency (**b, h**), percentage of silent neurons at resting membrane potential (**c, i**), resting membrane potential (**d, j**), input resistance (**e, k**), and voltage sag amplitude (**f, l**). **m**, Channelrhodopsin-assisted circuit mapping (CRACM) between Arc^AgRP^ and MPOA neurons. Whole-cell recordings from MPOA neurons, and 450 nm widefield stimulation of ChR2+ Arc^AgRP^ axons (see Methods). **n**, Example recording trace from MPOA neuron with sIPSCs. Note absence of light-evoked IPSCs in response to a train of 9 × 3-ms light pulses. **o**, Synaptic response pattern of MPOA neurons to acute Arc^AgRP^ terminal activation (11 cells from n = 3 mice). **p**, Representative example of action potentials in a ChR2-positive Arc^AgRP^ neuron evoked by single 3-ms light pulses. *U* test in **b, d**–**f, h, j**–**l.** Fisher’s exact test in **c, i**.

**Extended Data Fig. 8:**
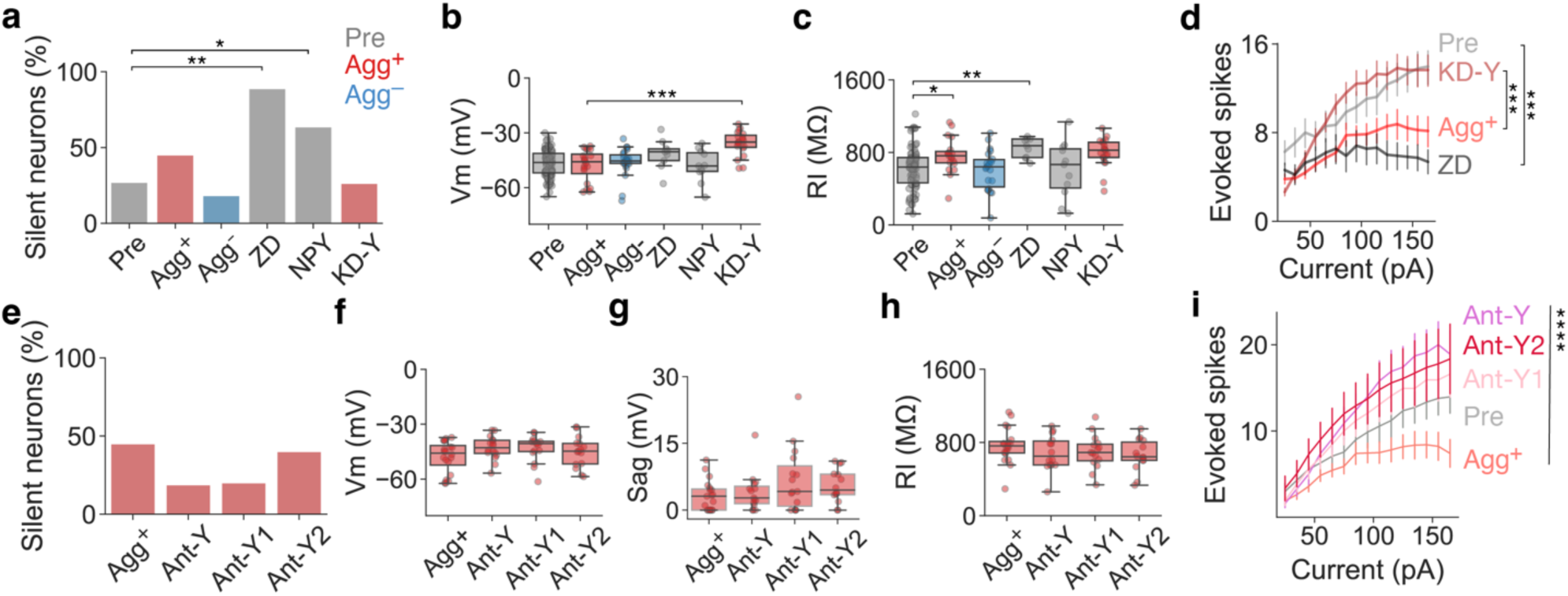
Biophysical effects of manipulating NPY signalling and HCN channel function. **a–c**, Biophysical parameters of MPOA neurons in mice before (Pre) and after (Agg^+^ and Agg^−^) 6-h food deprivation (82, 19, 22, 9, 11, 19 cells from n = 15, 6, 3, 1, 4, 3 mice): percentage of silent neurons at resting membrane potential (**a**), resting membrane potential (**b**), and input resistance (**c**). **d**, Action potentials per injected somatic current (Pre, Agg^+^, ZD, KD-Y; 10, 23, 17, 22 cells from n = 3, 7, 3, 5 mice). **e–i**, Biophysical parameters of MPOA neurons in brain slices of Agg^+^ mice after combined (Ant-Y) and separate (Ant-Y1, Ant-Y2) administration of Y1 and Y2 receptor antagonists (Pre, Agg^+^, Ant-Y, Ant-Y1, Ant-Y2; 19, 15, 15, 14, 12 cells from n = 6, 3, 2, 3, 3 mice): percentage of silent neurons at resting membrane potential (**e**), resting membrane potential (**f**), voltage sag amplitude (**g**), input resistance (**h**) and action potentials per injected current (**i**). Fisher’s exact test with Benjamini-Hochberg adjustment in **a, e**. *U* test with Benjamini-Hochberg adjustment in **b, c** (between Pre, Agg^+^ and Agg^−^; Pre, ZD and NPY; Agg^+^ and KD-Y). Two-way ANOVA with Tukey *post hoc* test in **d, i**. One-way ANOVA in **f, g, h**.

**Extended Data Fig. 9:**
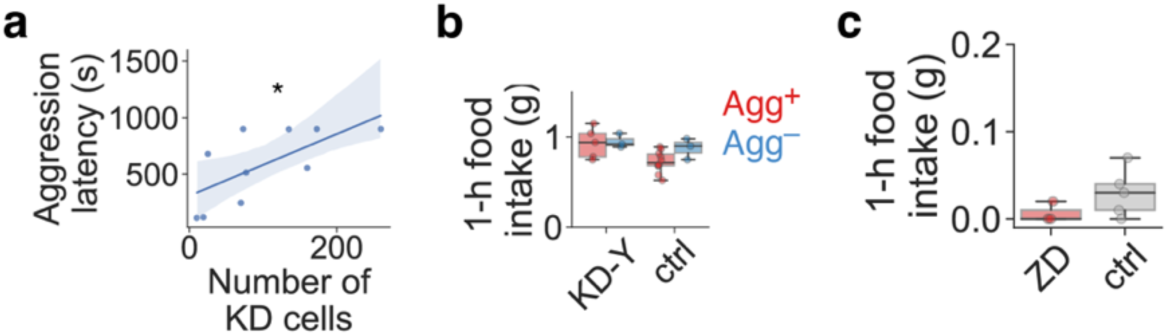
Behavioural effects of NPY knockdown and HCN blockade in the MPOA. a, Correlation between aggression latency and number of Arc^AgRP^→MPOA neurons transduced with KD-Y construct (linear regression, *r*^2^ = 0.450; *P* = 0.034; n = 10 mice). b, 1-h food intake of animals with knockdown of NPY in Arc^AgRP^→MPOA projection, and control (n = 8, 13 mice). c, 1-h food intake of animals with bilateral infusion of HCN blocker (ZD) or vehicle (ctrl) into MPOA (n = 3, 5 mice). *U* test in b, c.

**Extended Data Fig. 10:**
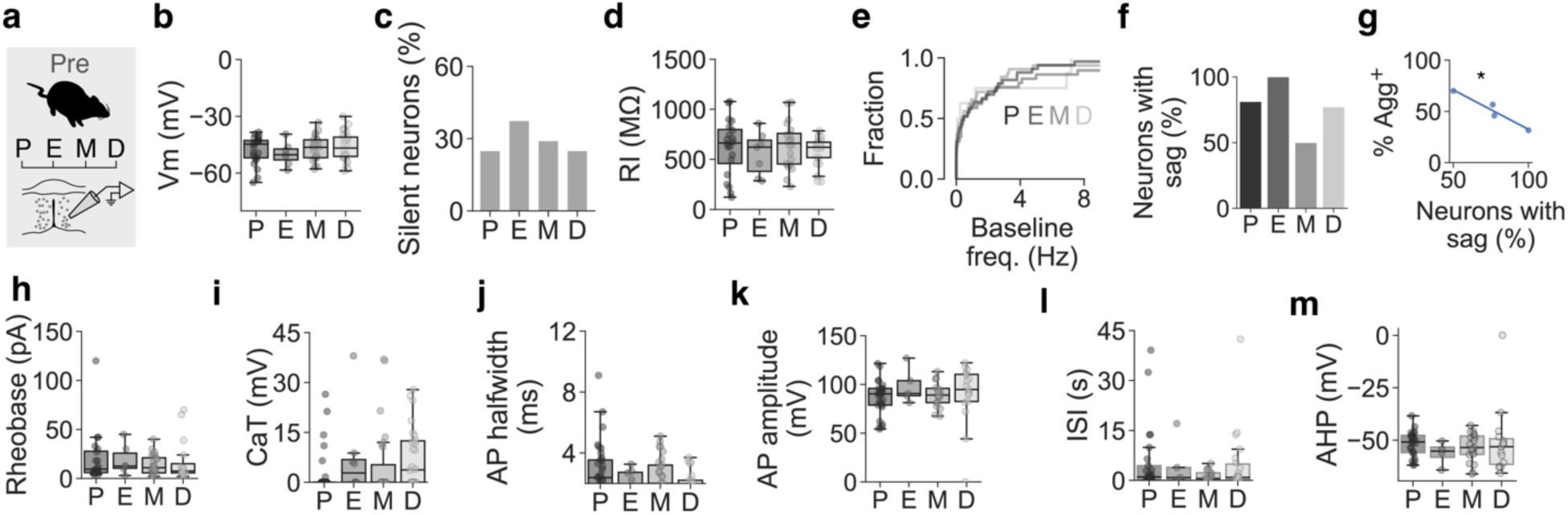
Biophysical effects of estrous state on MPOA neurons. **a–m**, Recordings from MPOA neurons at different estrous stages before food deprivation (**a**, P, proestrus, E, estrus, M, metestrus, D, diestrus; 31, 43, 31, 97 cells from n = 8, 8, 7, 7 mice): resting membrane potential (**b**), percentage of silent neurons at resting membrane potential (**c**), input resistance (**d**), baseline firing frequency (**e**), percentage of neurons with voltage sag (**f**), correlation between percentage of neurons with voltage sag and switching rate (**g**, linear regression, *r*^2^ = 0.99; *P* = 0.0059; data points are estrous stages), rheobase (**h**), rebound after hyperpolarisation due to T-type calcium currents (**i**), action potential half-width (**j**), action potential amplitude (**k**), inter-spike interval (**l**) and afterhyperpolarisation (**m**). One-way ANOVA in **b, d, e, h–m**. Fisher’s exact test in **c, f**.

**Extended Data Fig. 11:**
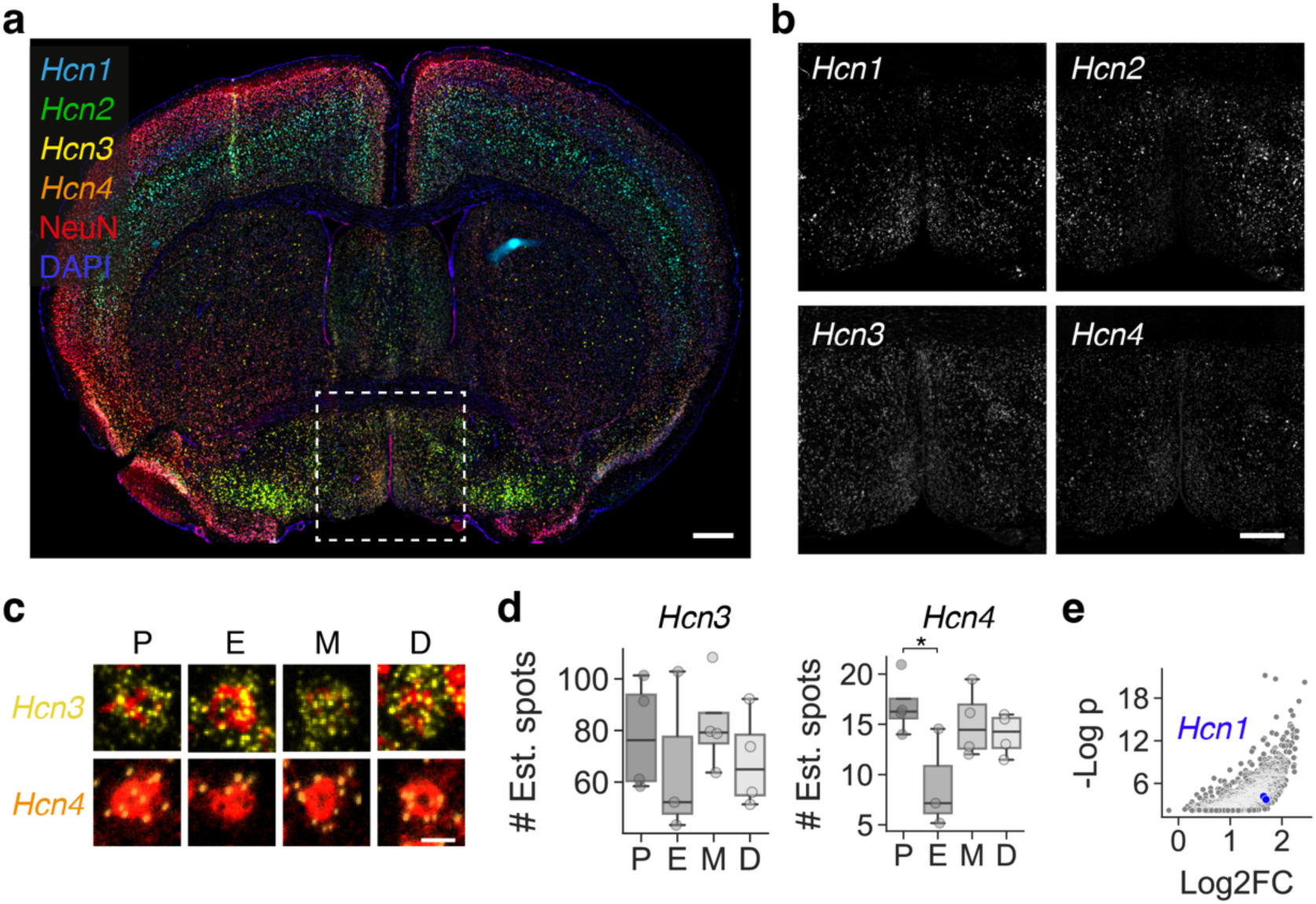
*Hcn* expression in the MPOA. **a**, Example coronal brain section with *Hcn* transcripts, after multiplexed *in situ* hybridisation and counterstaining with NeuN and DAPI. Scale bar, 500 µm. **b**, *Hcn* subunit expression in the MPOA. Scale bar, 300 µm. **c, d**, *Hcn3* and *Hcn4* mRNA expression across estrous cycle (**c**, red, NeuN counterstain; scale bars, 10 µm) and quantification (**d**, estimated number of spots, see Methods; n = 4, 3, 4, 4 mice). One-way ANOVA with Tukey *post hoc* test in **d**. **e**, RNA-seq of genes upregulated by E2 treatment in Esr1+ neurons across the mouse brain (data from ref. 66). p, *P* value. FC, fold change.

**Extended Data Fig. 12:**
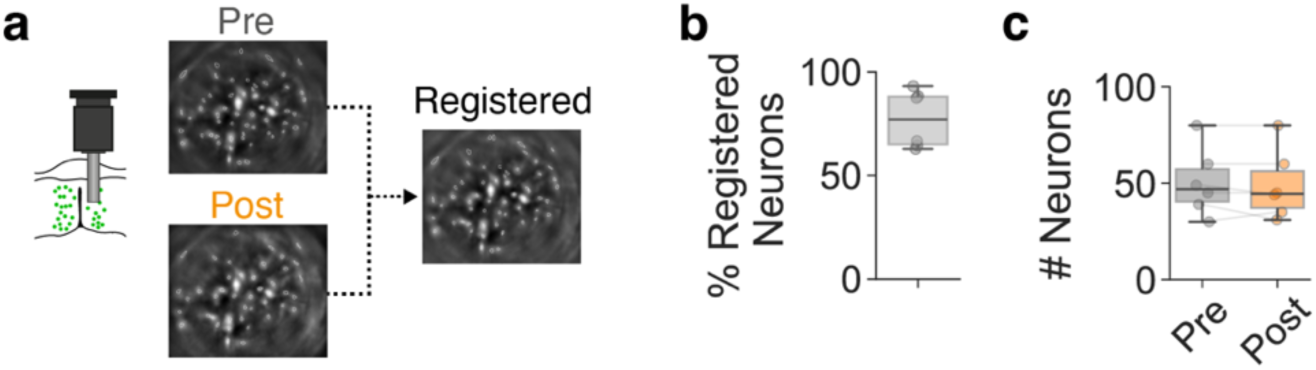
Longitudinal cell registration of micro-endoscopic images. a, Example and miniature microscope recording frames before and after registration (see Methods). b, Percentage of successfully registered neurons per animal (n = 5 mice). c, Number of detected neurons in Pre and Post recording sessions (n = 6 mice). *U* test in c.

**Extended Data Fig. 13:**
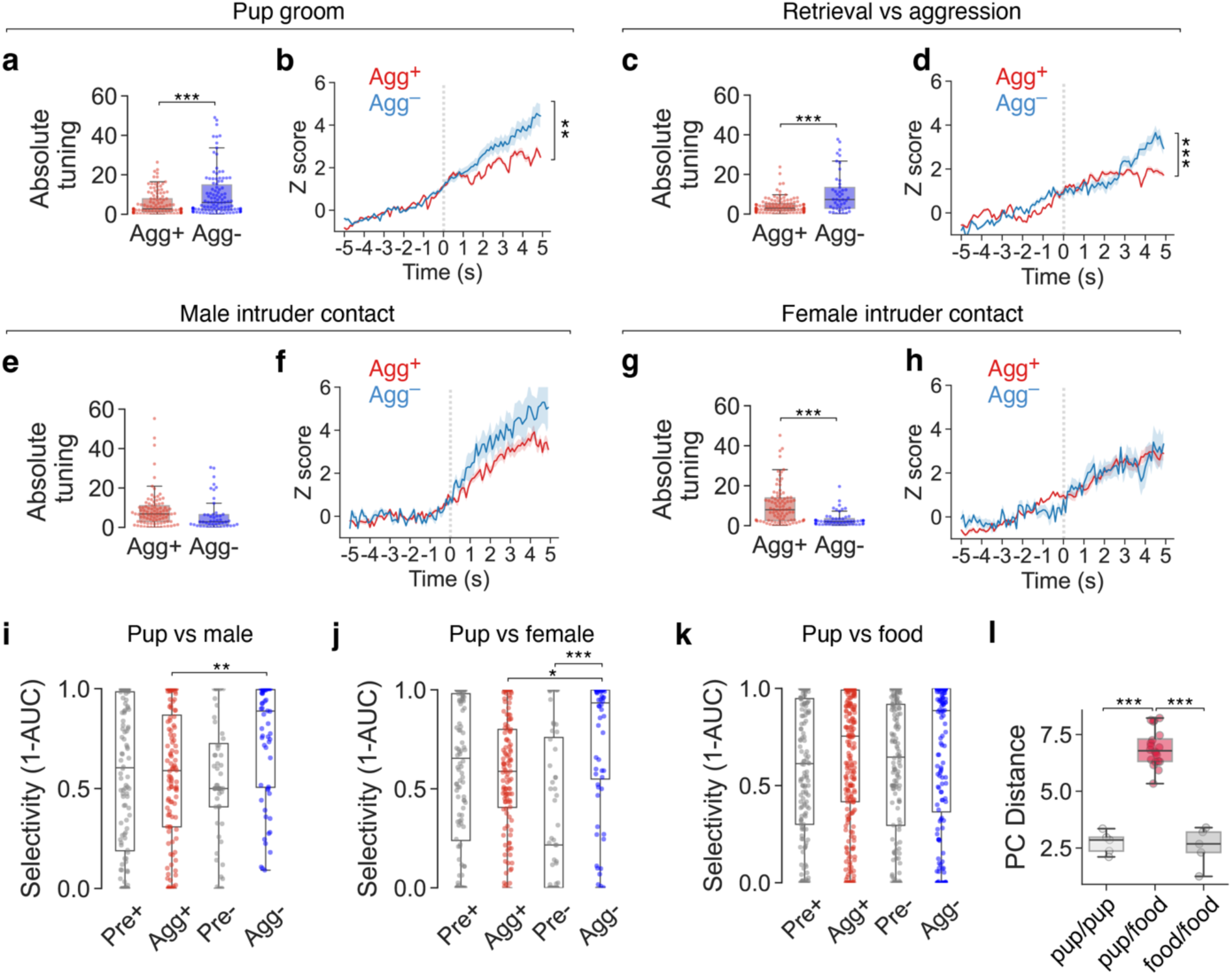
Enhanced MPOA neuronal responses to pup stimuli in Agg^−^mice. a–h, Averaged, *Z*-scored neuronal responses (a, c, e, g) and absolute tuning index (b, d, f, h) of MPOA neurons during indicated behaviours. i–k, Selectivity of chemoinvestigation-associated responses for indicated stimulus pairs compared with pups (116, 60 neurons from n = 3, 1 mice in i–j; 243, 148 neurons from n = 4, 2 mice in k). A selectivity score of 1 means the neuron is only activated during pup sniffing, a score of 0 means selective activation during sniffing of another stimulus, and a score of 0.5 is a nonselective response (see Methods). l, PC distance between Pre and Post episodes during pup vs food investigation (5, 20, 5 from 5 mice). Mixed linear model with mouse ID as group in a, c, e, g, i–k. One-way ANOVA with Tukey *post hoc* test in l. Two-way ANOVA with Tukey *post hoc* test in b, d, f, h.

**Extended Data Fig. 14:**
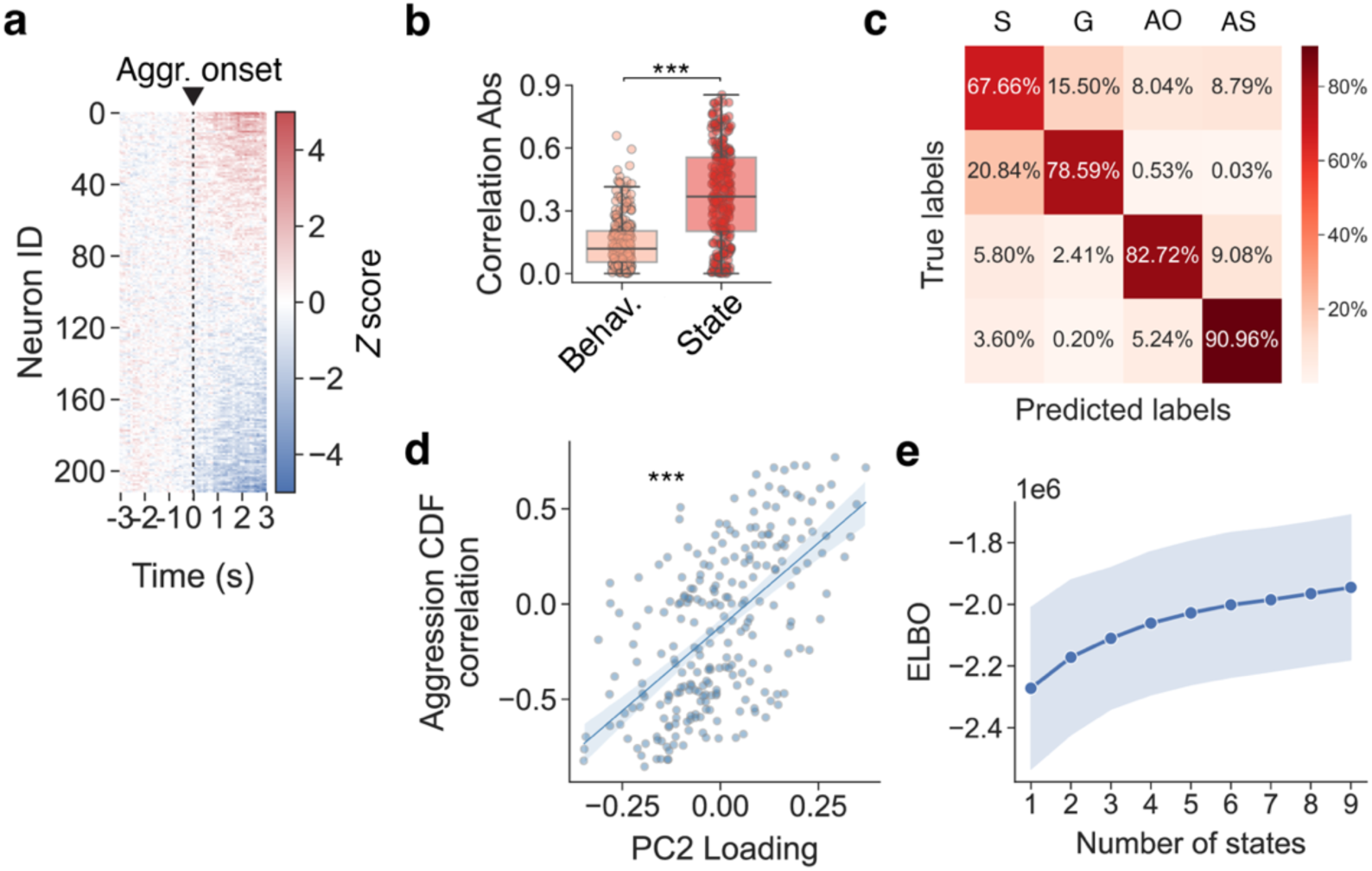
Aggression state encoding in the MPOA. a, Temporal profile of *Z*-scored neuronal responses during pup-directed aggression (average of aggression episodes 2–5) with hierarchical clustering based on mean response onset (243 neurons from n = 5 mice). b, Absolute correlation of MPOA neuronal activity with either aggression onset (‘Behaviour’) or with cumulative distribution function (CDF) of aggression state (n = 243, 243 neurons from 5 mice). c, Averaged confusion matrix from SVM trained on PC1 and PC2. S, pup sniffing, G, pup grooming, AO, first aggression, AS, later aggression. d, Correlation between each neuron’s loading on PC2 and its correlation coefficient with the CDF of aggression episodes (linear regression, *r*^2^ = 0.356, *P* = 4.46 × 10^25^, n = 243 from 5 mice). e, ELBO score for HMM fitting with variable numbers of states. Mixed linear model with mouse ID as group in b.

**Extended Data Fig 15:**
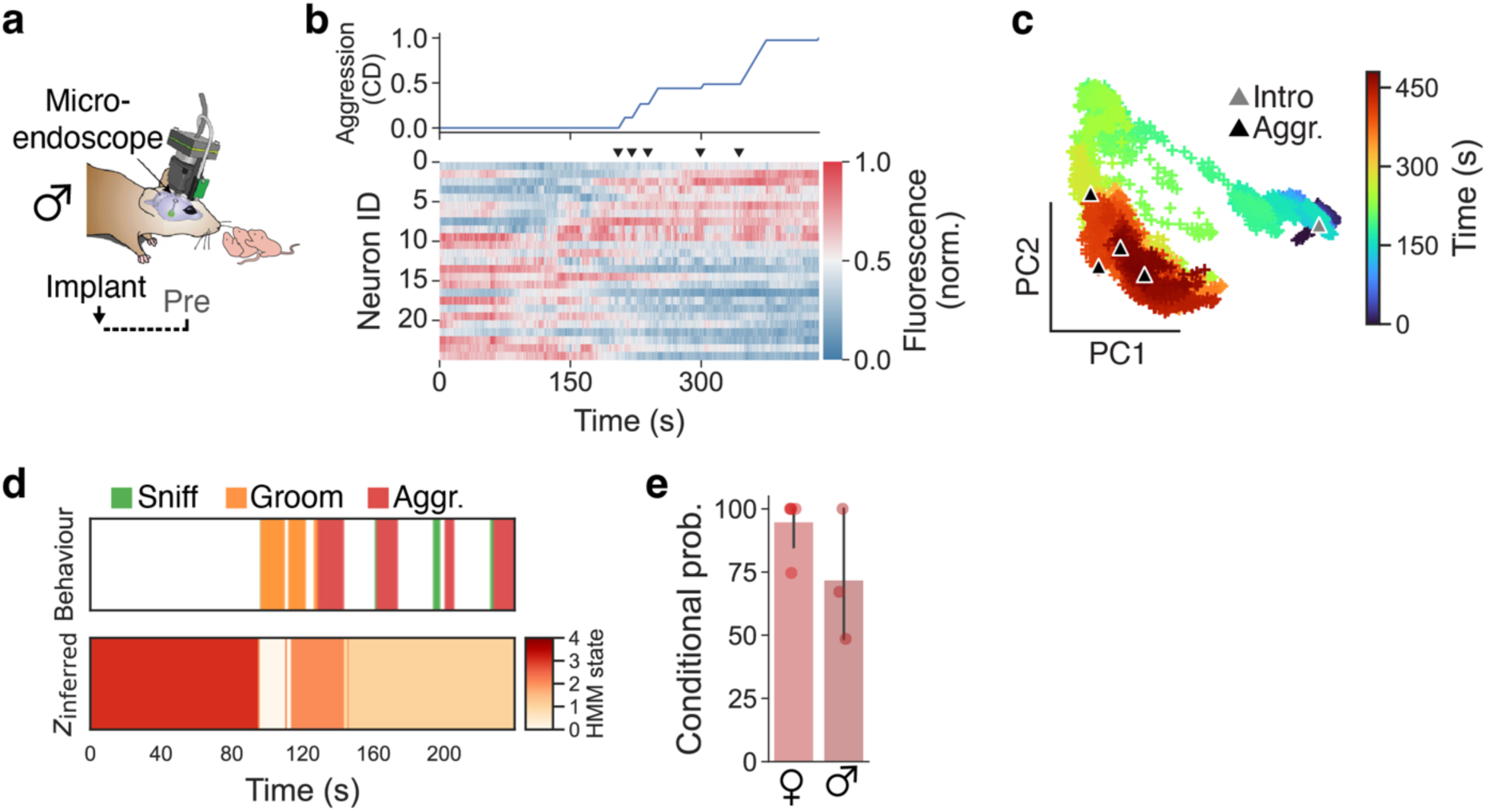
Aggression state encoding in the MPOA of male mice. **a**, Miniature microscope recordings during pup-directed aggression in males. **b**, Example heatmap of neuronal activity sorted based on correlation of activity with cumulative distribution (CD) of aggression (n = 25 neurons). Arrows indicate attack episodes. **c**, Population activity traces projected onto first two PCs. Note the aggression-specific state along PC2. **d**, Example HMM state segmentation using Agg^+^ neural data *(bottom)*, and ethogram *(top)*. **e**, Conditional probability of observing the indicated behaviour (aggression) when the system is in the Hidden Markov Model (HMM) state that most frequently aligns with aggression (females, males, n = 5, 3 mice).

**Extended Data Table 1:** Brain areas and coordinates. Injection, viral injection; stimulation, implantation coordinates for optic fibers (fiber tip position); recording, implantation coordinates for GRIN lenses or photometry fibers (tip position). Coordinates are AP (anteroposterior) / ML (mediolateral) / DV (dorsoventral) and in mm. DV coordinates are measured from brain surface.

| Acronym | Brain area | Injection | Stimulation | Recording |
| --- | --- | --- | --- | --- |
| AAA | Anterior amygdaloid area | – | – | – |
| ADP | Anterodorsal preoptic nucleus | – | – | – |
| AHA | Anterior hypothalamic area | – | – | – |
| ASO | Anterolateral striatal organ | – | – | – |
| ARC | Arcuate nucleus of hypothalamus | –1.5 / $\pm 0.25$ / –5.90 | – | –1.5 / 0.3 / –5.75 |
| AVP | Anteroventral preoptic nucleus | – | – | – |
| AVPV | Anteroventral periventricular nucleus | – | – | – |
| BLAa/p/v | Basolateral amygdaloid nucleus, anterior/posterior/ventral part | – | – | – |
| BNST | Bed nucleus of the stria terminalis | – | – | – |
| CEAc/l/m | Central amygdaloid nucleus, capsular/lateral/medial part | – | – | – |
| COAa/pl/pm | Cortical amygdaloid nucleus, anterior/posterolateral/posteromedial part | – | – | – |
| DMH | Dorsomedial hypothalamic nucleus | – | – | – |
| LA | Lateral amygdaloid nucleus | – | – | – |
| LH | Lateral hypothalamic area | – | – | – |
| LM | Lateral mammillary nucleus | – | – | – |
| LPOA | Lateral preoptic area | – | 0.0 / $\pm 1.0$ / –4.65 | – |
| LSc/r/v | Lateral septal nucleus, central/ventral part | – | – | – |
| MeA | Medial amygdaloid nucleus | – | – | – |
| MEPO | Median preoptic nucleus | – | – | – |
| MPO(A) | Medial preoptic area | 0.0 / $\pm 0.5$ / –5.05 | 0.0 / $\pm 0.5$ / –4.65 | 0.0 / 0.5 / –4.85 |
| MPN | Medial preoptic nucleus | – | – | – |
| MS | Medial septal nucleus | – | – | – |
| PA | Paraventricular thalamic nucleus, anterior part | – | – | – |
| PAG | Periaqueductal gray | – | – | – |
| PBN | Parabrachial nucleus | – | – | – |
| PD | Posterodorsal preoptic nucleus | – | – | – |
| PeF | Perifornical nucleus | – | – | – |
| PH | Posterior hypothalamic area | – | – | – |
| PMv | Ventral premammillary nucleus | – | – | – |
| PS | Parastrial nucleus | – | – | – |
| PVa/i/p/po | Periventricular hypothalamic nucleus, anterior/intermediate/posterior/preoptic part | – | – | – |
| PVH | Paraventricular hypothalamic nucleus | – | 0.7 / 0 /<br>–4.4 | – |
| PVT | Paraventricular nucleus of the thalamus | – | – | – |
| SCH | Suprachiasmatic nucleus | – | – | – |
| SUM | Supramammillary nucleus | – | – | – |
| TMd/v | Dorsal/ventral tuberomammillary nucleus | – | – | – |
| TU | Tuberal nucleus | – | – | – |
| VLPO | Ventrolateral preoptic nucleus | – | – | – |
| VMH | Ventromedial hypothalamic nucleus | – | – | – |
| VMPO | Ventromedial preoptic nucleus | – | – | – |
| VTA | Ventral tegmental area | – | – | – |

**Extended Data Table 2:** Blood plasma levels of P4 and E2 for estrous and pregnancy stages. Maximal values for each state are shown. P, proestrus, E, estrus, M, metestrus, D, diestrus, D10, pregnancy day 10, D18, pregnancy day 18.

| State | P4 plasma level<br>(ng / ml) | E2 plasma level<br>(pg / ml) | Ref. | $(P4 / E2) \times 10^{-3}$ | Switching rate |
| --- | --- | --- | --- | --- | --- |
| P | 64 | 10.3 | 105,106 | 6.21 | 0.57 |
| E | 25 | 4.2 | 105,106 | 0.95 | 0.31 |
| M | 27 | 3.1 | 105,106 | 8.71 | 0.7 |
| D | 15 | 7.5 | 105,106 | 2 | 0.46 |
| D10 | 37 | 16 | 46,47 | 2.31 | 0.44 |
| D18 | 70 | 6.8 | 46,47 | 1.03 | 0.33 |

## Supplementary Note

In a previous study, Li, et al. (2019) reported direct GABAergic connections from Arc^AgRP^ neurons onto 30% of MPOA neurons in female mice, using channelrhodopsin-assisted circuit mapping (CRACM)^34^. However, as the functional connectivity experiments in Fig. 2C of Li et al. (2019) were not performed with the sodium channel blocker tetrodotoxin (TTX) in the bath solution, it remains unclear whether the presented responses are monosynaptic. Indeed, the reported latencies of inhibitory postsynaptic currents (IPSCs) in females (mean ∼8 ms, Fig. 2D) do not, by themselves, support direct connectivity. In addition, the current traces in Fig. 2C of Li et al. (2019) indicate that the observed IPSCs are OFF-responses, i.e., occurring time-locked with the offset of optogenetic stimulation. Using CRACM with the same stimulation parameters as in Li et al. (2019), we were unable to detect direct GABAergic Arc^AgRP^→MPOA connections even though we could optogenetically evoke action potentials in ChR2-expressing Arc^AgRP^ cells in the same brain slices (Extended Data Fig. 7m–p). While we can only speculate about the reason(s) underlying this discrepancy, our findings do not support the existence of monosynaptic GABAergic Arc^AgRP^→MPOA connectivity.

